# Translational regulation of Pmt1 and Pmt2 by Bfr1 affects unfolded protein O-mannosylation

**DOI:** 10.1101/847095

**Authors:** Joan Castells-Ballester, Natalie Rinis, Ilgin Kotan, Lihi Gal, Daniela Bausewein, Ilia Kats, Ewa Zatorska, Günter Kramer, Bernd Bukau, Maya Schuldiner, Sabine Strahl

## Abstract

O-mannosylation is implicated in protein quality control in *Saccharomyces cerevisiae* due to the attachment of mannose to serine and threonine residues of un- or misfolded proteins in the endoplasmic reticulum (ER). This process also designated as unfolded protein O-mannosylation (UPOM) that ends futile folding cycles and saves cellular resources is mainly mediated by protein O-mannosyltransferases Pmt1 and Pmt2. Here we describe a genetic screen for factors that influence O-mannosylation in yeast, using slow-folding GFP as a reporter. Our screening identifies the RNA binding protein brefeldin A resistance factor 1 (Bfr1) that has not been linked to O-mannosylation and ER protein quality control before. We find that Bfr1 affects O-mannosylation through changes in Pmt1 and Pmt2 protein abundance, but has no effect on *PMT1* and *PMT2* transcript levels, mRNA localization to the ER membrane or protein stability. Ribosome profiling reveals that Bfr1 is a crucial factor for Pmt1 and Pmt2 translation thereby affecting unfolded protein O-mannosylation. Our results uncover a new level of regulation of protein quality control in the secretory pathway.

## INTRODUCTION

Glycosylation is a major protein modification that includes the addition of a sugar moiety onto a protein (Spiro, 2002). Two types of glycosylation conserved from fungi to humans are N-glycosylation and O-mannosylation (Neubert & Strahl, 2016). Both essential types of glycosylation start in the endoplasmic reticulum (ER) and share the common mannose donor Dol-P-mannose (Dol-P-Man). O-mannosylation entails the direct transfer of mannose from Dol-P-Man to serine and threonine residues of proteins entering the secretory pathway (herein referred to as secretory proteins) by different types of protein O-mannosyltransferase enzymes. Among those, only the protein O-mannosyltransferase (PMT) family is conserved among eukaryotes (Immervoll, Gentzsch, & Tanner, 1995; Jurado, Coloma, & Cruces, 1999; Lussier, Gentzsch, Sdicu, Bussey, & Tanner, 1995; Strahl-Bolsinger & Tanner, 1991; Willer, Amselgruber, Deutzmann, & Strahl, 2002). Changes in PMT-based O-mannosylation in humans result in genetic disorders called α-dystroglycanopathies (Brancaccio, 2019) and are also associated with various cancers (Carvalho, Reis, & Pinho, 2016; Kumari, Das, Adhya, Rath, & Mishra, 2019). In the baker’s yeast, *S. cerevisiae*, (from hereon termed simply yeast) O-mannosylation in the ER depends on PMTs only, making it an ideal model to study this crucial protein modification.

PMTs are ER membrane glycoproteins that have been shown to associate with the translocon to modify translocating polypeptides (Loibl et al., 2014). In yeast the redundant PMT family contains seven members, for six of which the O-mannosyltransferase activity has been proven. They are subdivided into three subfamilies referred to as PMT1 (Pmt1, Pmt5), PMT2 (Pmt2, Pmt3, Pmt6) and PMT4 (Pmt4) that show distinct substrate specificities (Loibl & Strahl, 2013). Pmt1-Pmt2 heterodimers contribute a major part of O-mannosyltransferase activity (Girrbach & Strahl, 2003).

Analysis of the yeast O-mannose glycoproteome revealed that around 20% of all ER and Golgi proteins are O-mannosylated many of those with crucial functions in protein glycosylation, folding, quality control and trafficking (Neubert et al., 2016). Hence it is not surprising that transcription of PMTs is enhanced under ER stress conditions (Travers et al., 2000) and general PMT inhibition induces the unfolded protein response (UPR) (Arroyo et al., 2011), a transcriptional response that regulates protein folding capacities of the ER and degradative processes termed ER associated degradation (ERAD) of un- or misfolded proteins (Hetz, 2012).

While most studies of O-mannosylation focus on the role of this modification during normal protein maturation along the secretory pathway, recently it has been demonstrated that there exists non-canonical O-mannosylation of proteins due to un- or misfolding (Xu & Ng, 2015a). This so-called unfolded protein O-mannosylation (UPOM) has been proposed as a molecular timer that is active in the early stages of ER protein quality control to abrogate futile folding cycles and save valuable cellular resources (Xu, Wang, Thibault, & Ng, 2013). Substrates that undergo UPOM have been shown to later be eliminated by the cell either by ERAD (Hirayama, Fujita, Yoko-o, & Jigami, 2008), vacuolar degradation (Coughlan, Walker, Cochran, Wittrup, & Brodsky, 2004) or cellular exclusion (Nakatsukasa et al., 2004). Similarly to O-mannosylation during maturation, modification during UPOM seems to also rely mostly on Pmt1 and Pmt2 (Goder & Melero, 2011; Xu et al., 2013). The most prominent UPOM substrate to date is slow-folding GFP that folds properly in the cytosol, but when targeted to the ER is recognized as a misfolded protein due to its slow folding and therefore gets O-mannosylated (Xu et al., 2013). O-mannosylation itself then blocks further folding of the fluorophore resulting in decreased fluorescence intensity rendering this protein an adequate reporter to monitor UPOM efficiency.

With the exception of Pmt1 and Pmt2 that mediate UPOM this protein quality control system is poorly defined. In the present study we screened for cellular factors that affect UPOM in yeast. To this end we took advantage of ER-targeted slow-folding GFP as a UPOM-reporter and identified brefeldin A resistance factor 1 (Bfr1) as an enhancer of Pmt1 and Pmt2 translation.

## RESULTS

### Genome-wide screen reveals Bfr1 as a factor influencing UPOM

To perform a genome-wide screen for identification of cellular factors affecting UPOM we took advantage of the model UPOM substrate, slow-folding ER-GFP (Xu et al., 2013). We stably introduced ER-GFP into the innocuous *HO* locus of *pmt1*Δ, *pmt2*Δ and *pmt4*Δ cells. ER targeting of GFP was ensured by an N-terminal Kar2 signal peptide and ER retention by a C-terminal HDEL retention signal (Fig. 1A, upper scheme). A fast folding variant of GFP (ER-GFPf) that escapes O-mannosylation and therefore changes in folding and fluorescence served as a negative control (Fisher & DeLisa, 2008). As shown in Fig. 1B ER-GFP shows reduced fluorescence compared to ER-GFPf expressed in wild type cells. In *pmt1*Δ and *pmt2*Δ cells reporter fluorescence is considerably enhanced compared to wild type whereas the GFP signal in *pmt4*Δ is not affected (Fig. 1B, C). These results are in line with previously published data in which ER-GFP is expressed from a centromeric plasmid (Xu et al., 2013). O-mannosylation of ER-GFP in wild type and *PMT* deletion mutants was monitored by probing lysates of respective cells for GFP (Fig. 1D). ER-GFP detection results in a main GFP signal accompanied by multiple higher molecular weight bands that are not seen in case of ER-GFPf (Fig. 1D, compare area designated by the white arrow in lanes 2 and 3). The same GFP pattern is detected in *PMT4* deficient (lane 6) but not *PMT1* and *PMT2* deficient cells (lanes 4 and 5) and correlates with O-mannosylation of ER-GFP. Treatment of immunopurified FLAG-tagged ER-GFP (Fig. 1A, lower scheme) with α1-2,3,6 mannosidase that removes O-linked α-mannose (Winterhalter, Lommel, Ruppert, & Strahl, 2013) confirmed that the signal above the main GFP band emanates from O-mannosyl glycans (Fig. 1E). We further examined whether ER-GFP expression that is driven by the strong *TDH3* promotor induces ER stress resulting in UPR induction (Fig. 1F). In contrast to ER-GFPf, expression of ER-GFP triggers the UPR as indicated by the significant increase of mRNA levels of the spliced (active) variant (Fig. 1F, *HAC1s*) of the UPR-inducing transcription factor Hac1 and the UPR-targeted Hsp70 chaperone Kar2. This suggests that at least in the case of GFP, slow folding rates rather than protein overexpression constitute the biggest challenge for the ER.

**Fig. 1.**
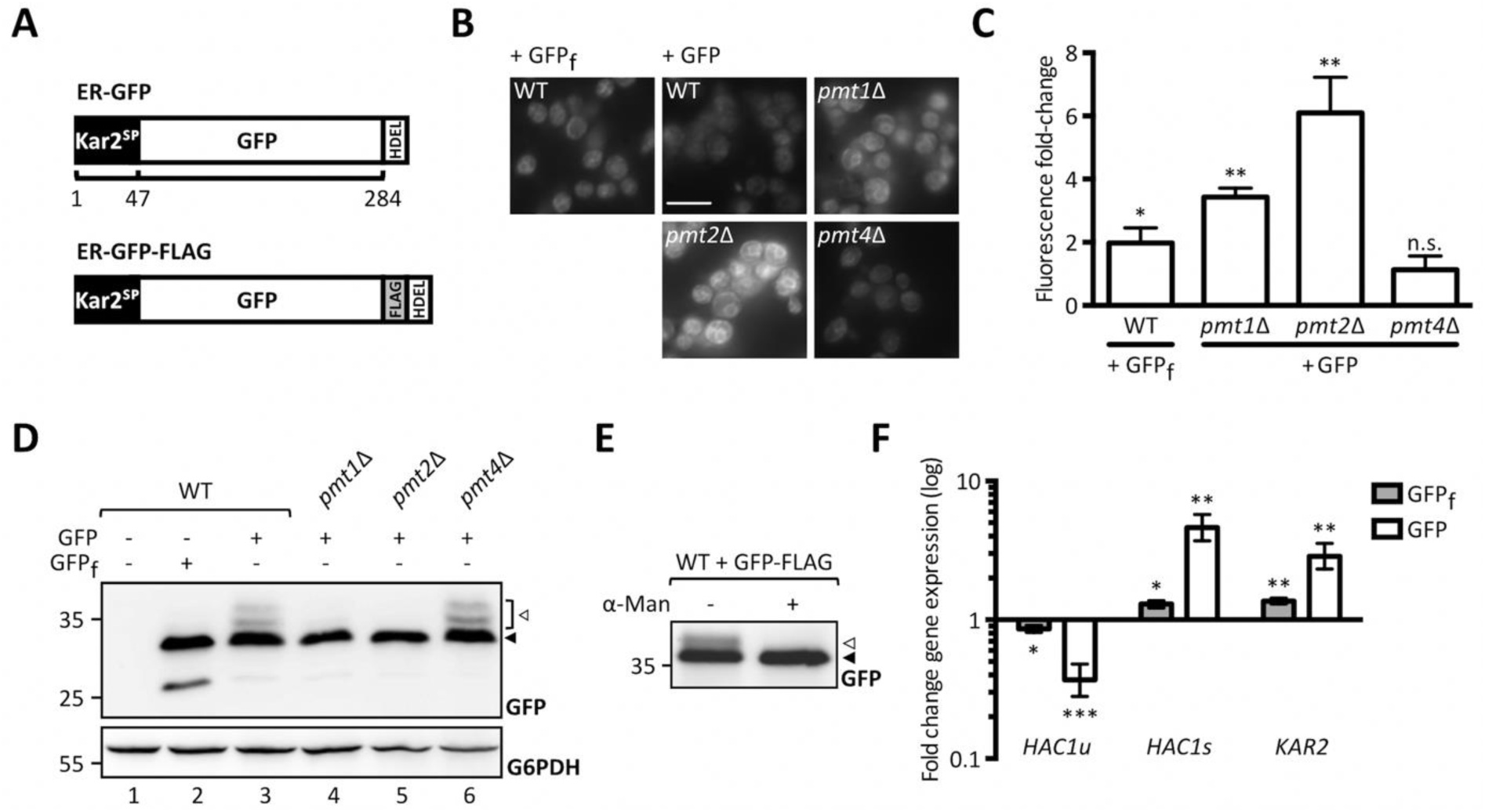
Analysis of ER-GFP as a UPOM-reporter. **A)** Schematic representation of ER-GFP N-terminally fused to the ER targeting signal peptide from Kar2 and C-terminally fused to the HDEL ER retention sequence (upper panel) and scheme of FLAG-tagged ER-GFP used for immunoprecipitation in C (lower panel). Fluorescence analysis of wild type and *pmtΔ* strains with genomically integrated ER-GFP by microscopy **(B)** and flow cytometry **(C)**. JEY06 (wild type ER-GFP), JCY010 (*pmt1*Δ ER-GFP), JCY011 (*pmt2*Δ ER-GFP), JCY012 (*pmt4*Δ ER-GFP) and JEY05 (wild type expressing ER-GFPf as negative control) cells were grown in YPD before being imaged under standard conditions (scale bar 10 μm) **(B)** or analyzed by flow cytometry **(C)**. In **(C)** fluorescent signals resulting from analysis of 20000 cells were normalized to wild type and results are plotted as fold-change. Error bars represent the range of values from three independent experiments. For statistical significance one-sample t-test was performed on log2(fold change). **D)** Western blot analysis of ER-GFP O-mannosylation in total cell extracts from strains shown in **(B)** and **(C)**. 20 μg of protein were resolved on a 12% PAA gel and detection was performed with an anti-GFP antibody. Wild type cells expressing ER-GFPf served as negative control and G6PDH was used as loading control. Arrows on the right indicate the main GFP signal (black arrow) and signals emanating from higher O-mannosylated GFP fractions (white arrow). **E)** FLAG-tag immunoprecipitation of ER-GFP on total cell extracts from wild type cells expressing FLAG-tagged ER-GFP from the pN014 plasmid. Purified ER-GFP-FLAG-HDEL was subjected to α1-2,3,6 mannosidase treatment overnight at 37°C and resolved on a 12% PAA gel. Detection was performed with an anti-GFP antibody. The signals emanating from higher O-mannosylated GFP-fractions (white arrow) collapse upon treatment into the main GFP signal (black arrow). Results are representative of two independent experiments. **F)** RT-PCR analysis of *HAC1u*, *HAC1s* and *KAR2* mRNA levels in wild type cells expressing ER-GFPf and ER-GFP respectively. JEY05 (wild type ER-GFPf) and JEY06 (wild type ER-GFP) cells were grown in YPD, total RNA was extracted, and cDNA was prepared and used as a template for RT-PCR. Fold-change was calculated from three independent experiments with respect to *ACT1* mRNA and error bars represent the confidence interval. For statistical significance one-sample t-test was performed on log2^−ΔΔCt^. N.s.=not significant

As depicted in Fig. 2A, the ER-GFP expressing wild type strain was crossed with libraries containing viable deletion strains of non-essential genes and hypomorphic mutants of essential ones to create new libraries in which each haploid strain expresses the ER-GFP on the background of one mutant allele. The median fluorescence intensities (MFIs) of all viable strains resulting upon crossing are shown in Fig. 2B (small diagram on the right) and a detailed listing of all identified targets is available in Suppl. Table S1. Analysis of ER-GFP median intensity frequency distribution for more than 5000 viable mutant strains revealed that approximately 5% displayed fluorescence exceeding the MFI range of ER-GFP in wild type cells (Fig. 2B, zoomed in area and green bars in bar diagram). A total of 109 genes exceeded the threshold (median GFP intensity at 187, red dotted line in Fig. 2B) and were considered as positive hits (Suppl. Table S1). Validity of the screen was confirmed by the presence of *PMT1* (position 38) and *PMT2* (position 3) among the positive candidates. Further analysis of screening hits was performed by manual assessment of GFP signal localization to the ER. Out of 109 candidates, only *spf1*Δ cells showed predominant cytosolic GFP fluorescence further confirmed in an independent *spf1*Δ mutant by fluorescence microscopy (Suppl. Fig. 1A). Among the residual 108 candidates stress pathway components (e.g. *oca1*Δ and *oca2*Δ involved in oxidative stress response; *sln1*Δ, *ptc1*Δ and *sic1*Δ encoding for functional components of the high osmolarity glycerol (HOG) pathway) and components of N-glycosylation and quality control (e.g. *ost3*Δ (Suppl. Fig. 2) and *cwh41*Δ) were present. Analysis of the O-mannosylation status of the canonical Pmt1-Pmt2 client Hsp150 revealed that the vast majority of the mutants do not severely affect O-mannosylation in general, judging by the prevalence of the molecular mass of Hsp150 upon the gene deletions. However, in a substantial number of mutants, we observed the presence of subspecies of Hsp150 that likely result from general maturation defects (Suppl. Table S1). Among those are for example *ost3*Δ and *pop2*Δ (Suppl. Fig. 1B) that affect N-glycosylation and mRNA catabolism, respectively, and for which general defects in protein homeostasis have been reported previously (Preissler et al., 2015; Stevens et al., 2017). Since we were especially interested in candidates that directly affect glycosylation of the UPOM-reporter systematic analysis of candidate genes was performed by determining ER-GFP O-mannosylation by Western blot. This revealed that for most of the tested mutant strains increased GFP fluorescence did not correlate with significantly reduced O-mannosylation (Suppl. Table S1). Next to *pmt1*Δ and *pmt2*Δ, only two additional mutants were found to abrogate ER-GFP O-mannosylation: *bfr1*Δ (Fig. 2C, D) and *psa1^DAmP^* (Suppl. Fig. 3A). *PSA1* is an essential gene encoding for the enzyme GDP-mannose pyrophosphorylase that is responsible for the synthesis of GDP-mannose, the mannose donor in Dol-P-Man synthesis (Hashimoto, Sakakibara, Yamasaki, & Yoda, 1997) (Suppl. Fig. 3B). Since decreased expression of Psa1 in the *psa1^DAmP^* most likely limits availability of the mannose donor Dol-P-Man thereby affecting PMT activity, we decided to herein focus on *BFR1* whose role remains unknown.

**Fig. 2.**
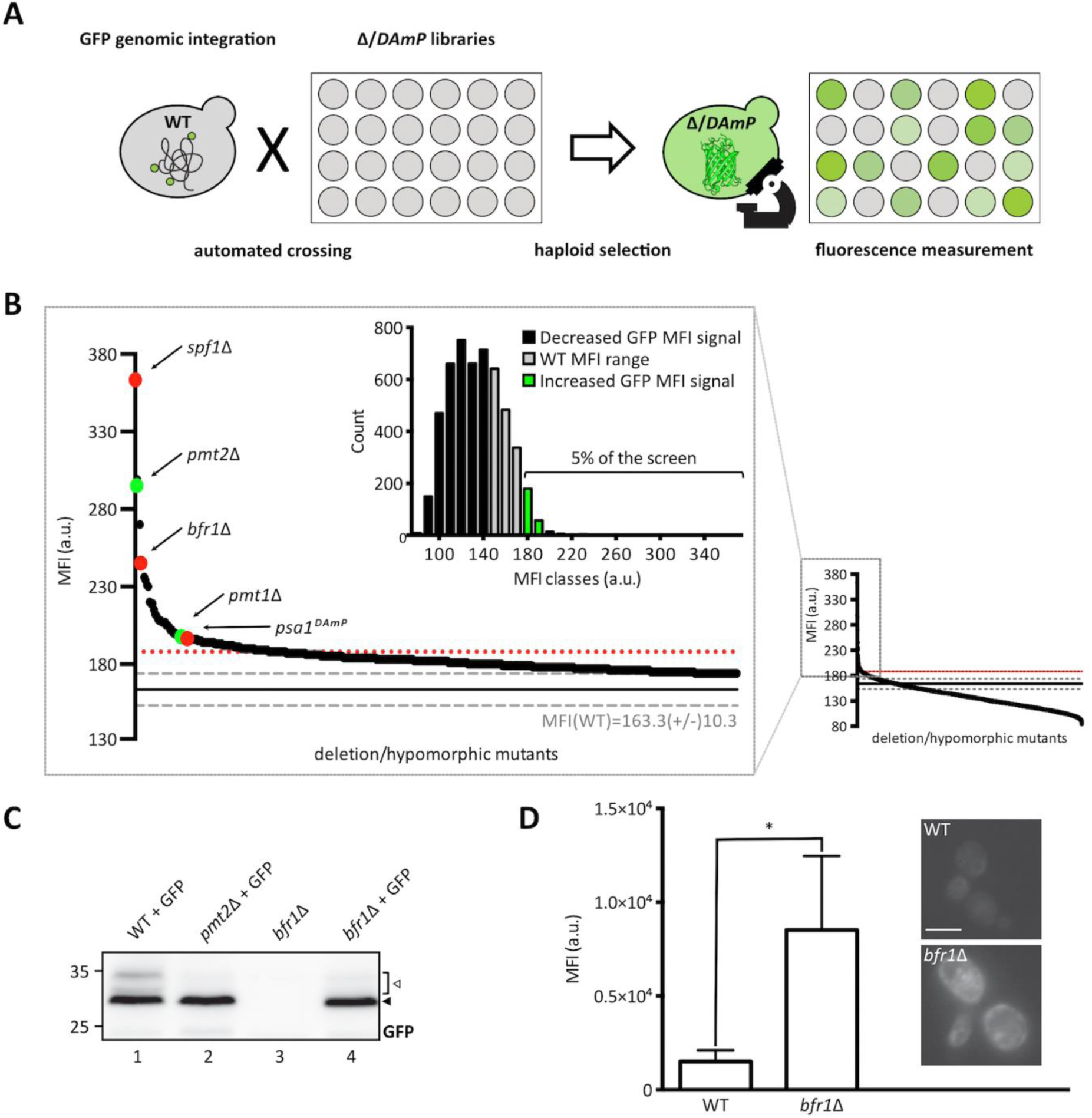
Identification of Bfr1 in a genome-wide UPOM screen. **A)** Schematic flowchart representing the major steps of the genome-wide high-throughput screen for identification of UPOM factors using ER-GFP as a fluorescent reporter. In brief, the ER-GFP expressing JEY06 strain was crossed with the yeast deletion library (Giaever et al., 2002) and the DAmP library (Breslow et al., 2008) on 1536 colony format YPD plates. Obtained diploids were selected for ER-GFP as well as deletion/DAmP mutations using KanR and URA3 respectively. Sporulation was induced upon nitrogen starvation for 7 days and haploid cells were selected on SD plates with aforementioned selections as well as toxic amino acid derivatives to eliminate residual diploids. Haploids were immobilized on Concanavalin A coated 384 well format microscopy plates and analyzed using an automated microscopy setup (Breker et al., 2013). **B)** Graphic representation of screening results. The small graph on the right represents the median fluorescence intensity (MFI) distribution of mutant strains (x-axis) analyzed within the UPOM screen. The magnified region on the left contains mutants that display fluorescence intensities that exceed the range of wild type MFI (163.6 +/− 10.3, indicated by grey dashed lines). The threshold for positive hit selection is marked by a red dotted line. Coloured dots depict expected hits (*PMT1* and *PMT2* in green) and hits that were further analyzed (*SPF1*, *BFR1* and *PSA1* in red). The bar graph on the upper left represents the median intensity frequency distribution of all mutant strains. Bars in grey depict frequencies of mutant strains with fluorescence within the wild type MFI range. Black and green bars represent frequencies of mutant strains with GFP signals below or above the wild type MFI range respectively. **C)** Western blot analysis of ER-GFP O-mannosylation in total cell extracts from wild type (BY4741) and *bfr1*Δ strains from the ER-GFP screen. Equivalents to 0.2 OD600 were resolved on a 12% PAA gel and detection was performed with an anti-GFP antibody. A *pmt2*Δ strain expressing ER-GFP served as a positive control. Arrows on the right indicate the main GFP signal (black arrow) and signals emanating from higher O-mannosylated GFP fractions (white arrow). In *bfr1*Δ cells no higher O-mannosylated GFP fractions are detected. **D)** JEY06 (wild type) and a *de novo* generated *bfr1*Δ strain in which ER-GFP was genomically integrated were grown in YPD media before being analyzed by flow cytometry or imaged under standard conditions (scale bar 5 μm). Fluorescent signal resulting from analysis of 20000 cells via flow cytometry was normalized to wild type and error bars represent the range of values from three independent experiments. For statistical significance one-sample t-test was performed on log2(fold change).

We verified that in *bfr1Δ* cells, a strong decrease of ER-GFP O-mannosylation (Fig. 2C, compare lanes 1 and 4) goes hand in hand with significantly increased ER-GFP fluorescence compared to wild type as assessed by flow cytometry and fluorescence microscopy (Fig. 2D), in agreement with improved folding of the reporter. Enhanced ER-GFP fluorescence upon *BFR1* deletion was confirmed via flow cytometry in several independent mutants (Suppl. Fig. 4). All in all, our screen uncovered an unexpected role for Bfr1 in O-mannosylation.

### Bfr1 affects UPOM by modulating Pmt1 and Pmt2 protein levels

*BFR1* was identified in a genetic screen as a multicopy suppressor of brefeldin A induced lethality in yeast (Jackson & Kepes, 1994). It is associated with mRNA metabolism as it was shown to interact with the RNA binding protein Scp160 in polyribosome associated mRNP complexes (Lang, Li, Black-Brewster, & Fridovich-Keil, 2001). Since then mRNA related functions of Bfr1 have gained increasing attention: Bfr1 was shown to affect P-body formation (Simpson, Lui, Kershaw, Sims, & Ashe, 2014; Weidner, Wang, Prescianotto-Baschong, Estrada, & Spang, 2014) and to bind hundreds of mRNAs despite the fact that it lacks canonical RNA binding domains (Hogan, Riordan, Gerber, Herschlag, & Brown, 2008; Lapointe, Wilinski, Saunders, & Wickens, 2015).

Considering the role of Bfr1 in mRNA metabolism and the recent finding that Bfr1 binds *PMT1* and *PMT2* transcripts (Lapointe et al., 2015) we hypothesized that Bfr1 could affect UPOM by modulating Pmt1 and Pmt2 protein levels. We therefore analyzed Pmt1 and Pmt2 protein abundance in wild type versus *bfr1*Δ cells (Fig. 3A, B, left panels). Our results show that Pmt1 and Pmt2 protein levels are markedly reduced in *BFR1* deficient versus wild type cells. This holds true under ER stress conditions caused by the ER-GFP reporter in the screening strain background (compare lanes 2 and 4 in Fig. 3A, B) with Pmt1 and Pmt2 levels increased in response to UPR (compare lanes 1 and 2 in Fig. 3A, B) as well as in the absence of ER-GFP in an independent strain background (compare lanes 1 and 3 in Fig. 3A, B). Quantification of PMT protein levels reveals a significant 2-fold reduction for both PMTs (Fig. 3A, B, right panels) in *bfr1*Δ versus wild type cells.

**Fig. 3.**
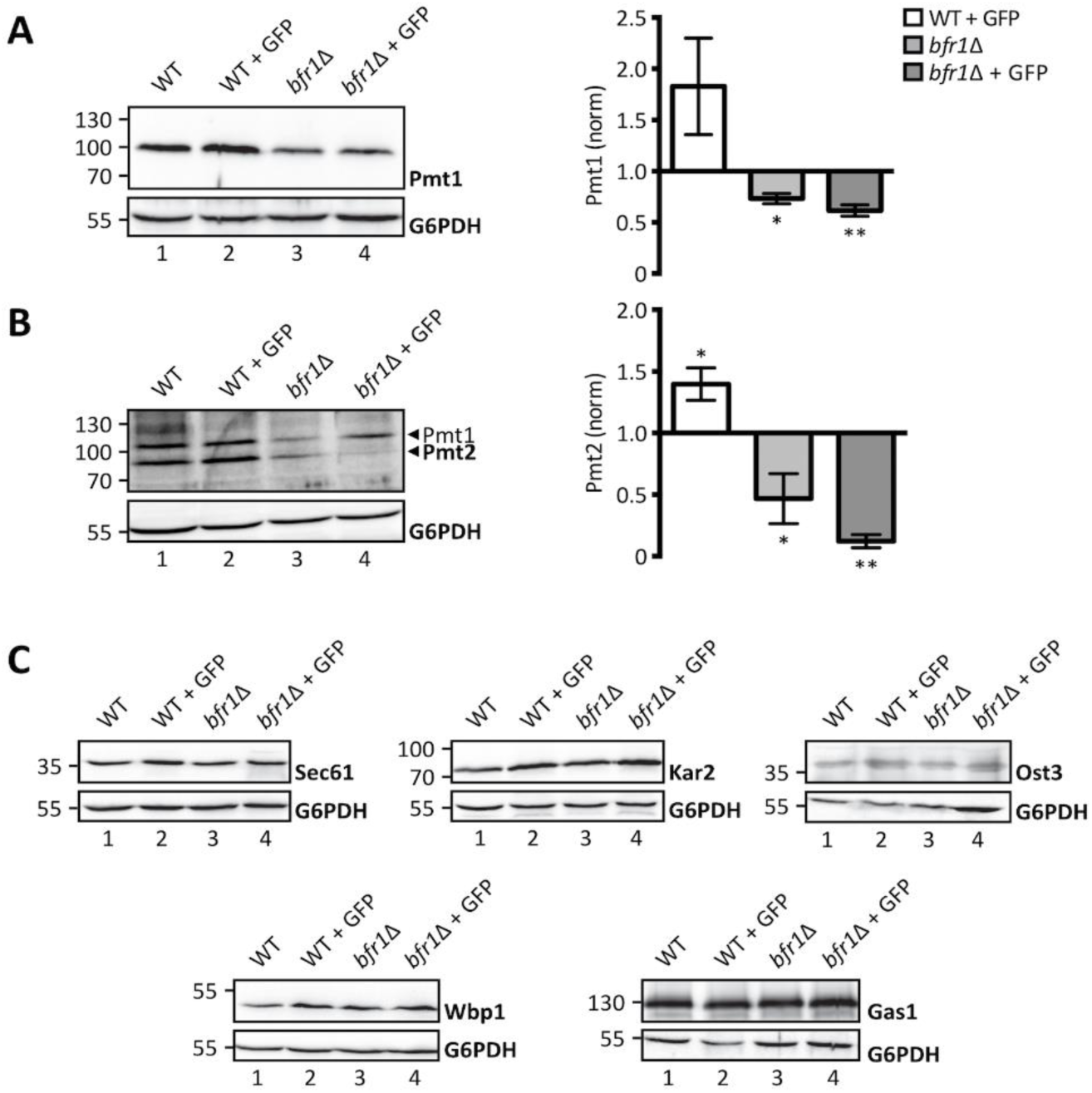
Pmt1 and Pmt2 protein levels are reduced in *BFR1* deficient cells. Western blot analysis of protein levels of Pmt1 **(A)**,Pmt2 **(B)**, and other representative proteins known to be targeted by Bfr1 (Lapointe et al., 2015) **(C)** in total cell extracts from wild type (BY4741), JEY06 (wild type ER-GFP), *bfr1*Δ and *bfr1*Δ ER-GFP strains. **(A)** and **(B)** left panels: 20 μg of protein were resolved on a 12% PAA gel and detection was performed with an anti-Pmt1 and anti-Pmt2 antibody respectively. Pmt2 detection was initially performed with a polyclonal serum detecting Pmt1 at the same time (lower and upper band indicated by black arrows respectively). In subsequent experiments preabsorption of the polyclonal serum was performed on membranes from *PMT2* deficient cells (single band detection for Pmt2 in e.g. Fig. 4). G6PDH was used as loading control. **(A)** and **(B)** right panels: Western blot signals were quantified using Image Studio Lite v 5.2 and PMT signals were first normalized to the respective G6PDH signals and subsequently normalized to the Pmt/G6PDH ratio calculated for wild type cells. Error bars represent the range of values from three independent experiments. For statistical significance one-sample t-test was performed on log2(fold change). **C)** 20 μg of protein were resolved on a 12% PAA gel and detection was performed with the indicated antibodies. G6PDH was used as loading control and results are representative of three independent experiments.

Since Bfr1 binds to numerous mRNAs we investigated the effect of *BFR1* deletion on protein levels of representative Bfr1 interactors (Lapointe et al., 2015) involved in protein import such as the main translocon subunit Sec61 (Deshaies & Schekman, 1987), quality control such as the Hsp70 chaperone Kar2 (Rose, Misra, & Vogel, 1989) and N-glycosylation such as oligosaccharyl transferase (OST) subunits Ost3 and Wbp1 (Karaoglu, Kelleher, & Gilmore, 1995; te Heesen, Janetzky, Lehle, & Aebi, 1992). We also analyzed protein levels of the GPI-anchored protein Gas1 that is highly O-mannosylated (Nuoffer, Jeno, Conzelmann, & Riezman, 1991) (Fig. 3C). Results show no major changes in protein levels for any of these Bfr1 targets in wild type versus *bfr1*Δ cells (compare lane 1 with 3 and 2 with 4 in Fig. 3C) suggesting that Bfr1 binding to mRNA alone is not sufficient to affect protein abundance.

To further substantiate the finding that O-mannosylation defects observed upon *BFR1* deletion result directly from decreased protein levels of Pmt1 and Pmt2, we performed a functional rescue experiment by overexpressing Pmt2. As shown in Fig. 4A Pmt2 overexpression restores O-mannosylation of ER-GFP in *BFR1* deficient cells. In agreement Pmt2 overexpression significantly reduces GFP fluorescence detected in *bfr1*Δ cells, however, not to wild type levels (Fig. 4B). In *BFR1* deficient cells Pmt2 protein levels are markedly decreased compared to wild type (Fig. 4C, compare lanes 1 and 3), even upon Pmt2 overexpression (compare lanes 2 and 4) and irrespective of ER stress caused by ER-GFP expression (compare lane 5 with 7 and 6 with 8). Inability to restore native Pmt2 levels as well as reduced levels of Pmt1 may explain why full complementation of ER-GFP O-mannosylation could not be gained. Taken together, our data show that the aberrant O-mannosylation of ER-GFP in *bfr1*Δ cells is a direct consequence of decreased Pmt1 and Pmt2 protein levels and that Bfr1 affects UPOM by controlling the abundance of these enzymes.

**Fig. 4.**
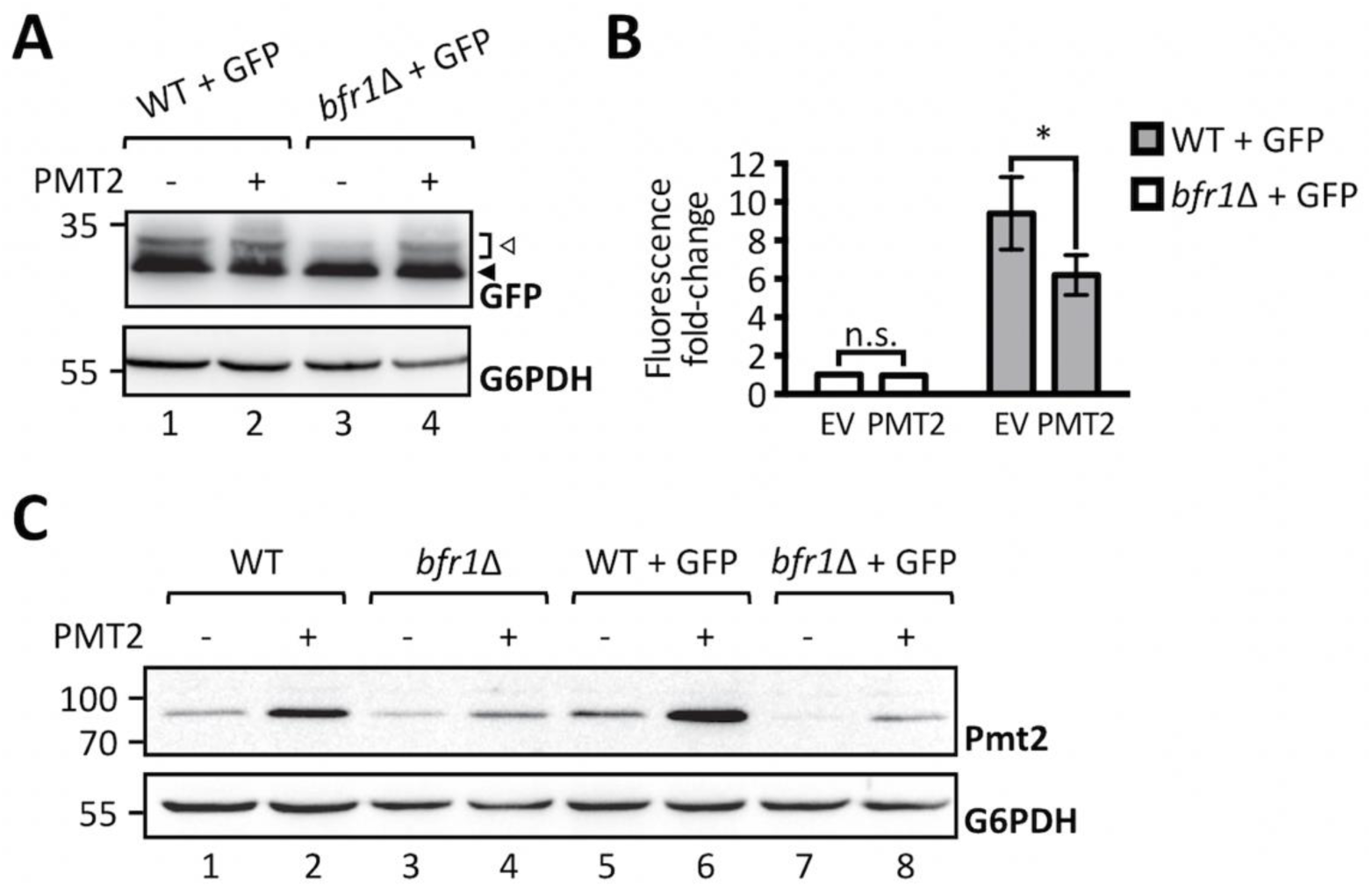
Pmt2 overexpression partially rescues loss of ER-GFP O-mannosylation in *bfr1*Δ cells. **A)** Western blot analysis of ER-GFP O-mannosylation in total cell extracts from JEY06 (wild type ER-GFP) and *bfr1*Δ ER-GFP strains transformed with either pRS41N (empty vector) or pJC09 (*PMT2*). Strains were grown under standard conditions in YPD supplemented with nourseothricin for selection. 20 µg of protein were resolved on a 12% PAA gel and detection was performed with an anti-GFP antibody. G6PDH was used as loading control. Arrows on the right indicate the main GFP signal (black arrow) and signals emanating from higher O-mannosylated GFP fractions (white arrow). Pmt2 overexpression partially restores ER-GFP O-mannosylation in the *bfr1*Δ strain. **B)** Flow cytometry analysis of strains described in **(A)**. Fluorescent signal for each strain resulted from analysis of 20000 cells and statistical significance was assessed by a 2way ANOVA on three independent experiments. Pmt2 overexpression partially restores ER-GFP fluorescence to the level detected in the JEY06 strain. **C)** Western blot analysis of Pmt2 protein levels in total cell extracts from wild type (BY4741), *bfr1*Δ, JEY06 (wild type ER-GFP) and *bfr1*Δ ER-GFP strains transformed with either pRS41N (empty vector) or pJC09 (*PMT2*) and grown as in **(A)**. 20 µg of protein were resolved on a 12% PAA gel and detection was performed with an anti-Pmt2 antibody. G6PDH was used as loading control. N.s.=not significant

### Bfr1 affects Pmt1 and Pmt2 protein levels on a posttranscriptional level

We next analyzed whether Bfr1 affects *PMT* transcript levels by measuring *PMT1* and *PMT2* mRNA abundance in wild type versus *bfr1*Δ cells (Fig. 5A). No significant changes in mRNA levels for both PMTs were found. To exclude impact of mRNA 5’regions *PMT2* was placed under the control of a *GAL1* inducible promotor and Pmt2 protein as well as transcript levels were assessed in wild type versus mutant cells. Pmt2 protein levels were markedly reduced in *BFR1* deficient cells (Fig. 5B, compare lanes 3 and 4) whereas transcript levels were unaffected (Fig. 5C). These results pointed to a posttranscriptional control of PMT synthesis mediated by either reduced translation or reduced protein stability in *bfr1*Δ cells. Cycloheximide chase experiments demonstrated that protein stability was not affected in *bfr1*Δ mutants (Fig. 5D), suggesting a possible effect of Bfr1 on translation efficiency.

**Fig. 5.**
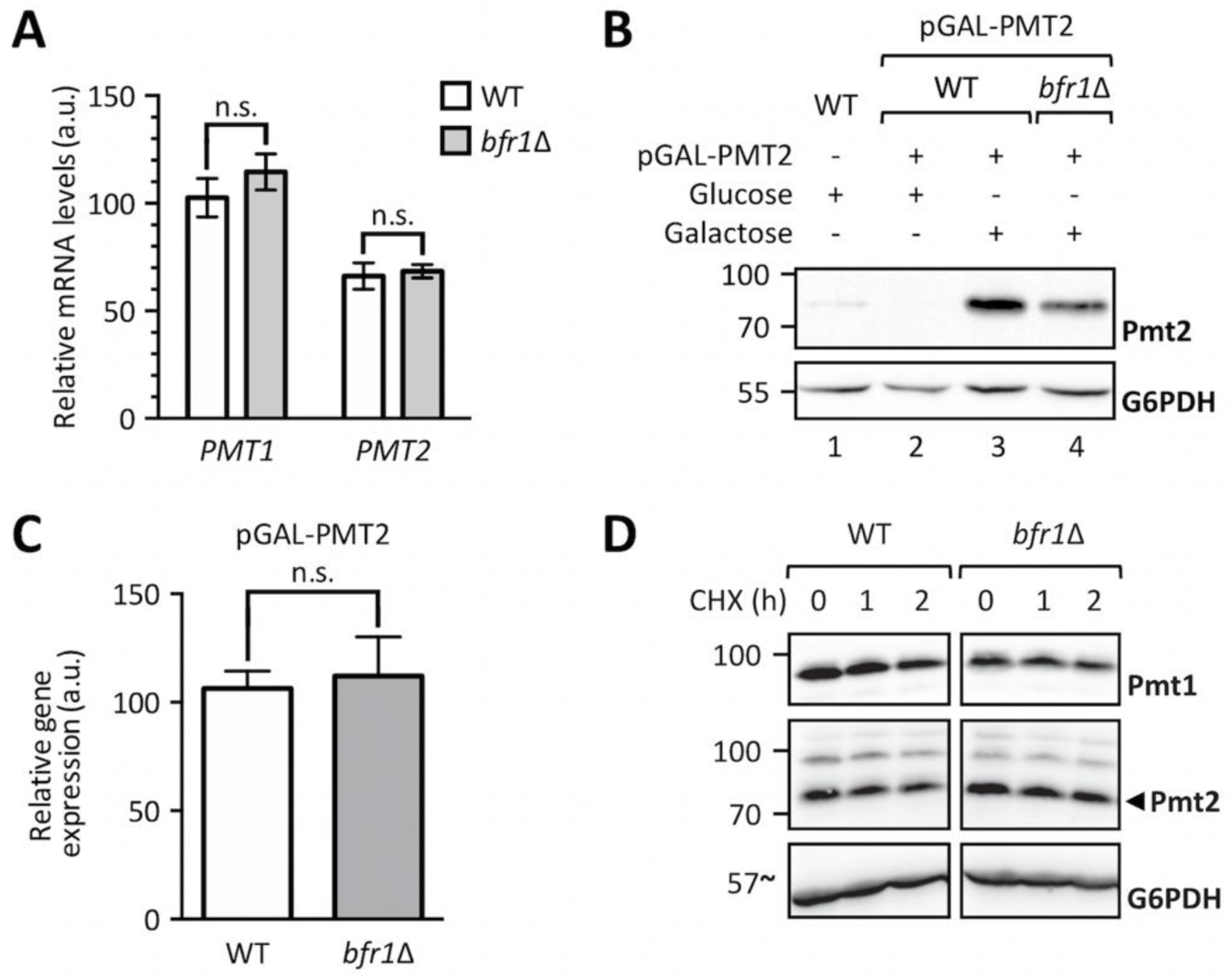
*BFR1* deletion does not affect *PMT1* and *PMT2* transcription. **A)** RT-PCR analysis of *PMT1* and *PMT2* mRNA levels in wild type (BY4741) and *bfr1*Δ strains. Cells were grown in YPD medium, total RNA was extracted and cDNA was prepared and used as a template for RT-PCR. Results show mRNA abundance with respect to *TAF10* mRNA from three independent experiments ± SD. For statistical significance a multiple t-test was performed. **B)** Western blot analysis of Pmt2 protein levels expressed under the control of a *GAL1* inducible promotor in total cell extracts from strains described in **(A)** upon addition of indicated sugars. **C)** RT-PCR analysis of *PMT2* mRNA levels in strains described in **(A)** in which Pmt2 is expressed under the control of a *GAL1* inducible promotor upon growth in galactose containing media. Results show mRNA abundance with respect to *ACT1* mRNA from three independent experiments ± SD. For statistical significance an unpaired t-test was performed. N.s.=not significant

### *BFR1* deletion does not affect *PMT1* and *PMT2* mRNA localization to the ER

Cotranslational protein translocation requires targeting of mRNAs encoding for secretory proteins to the ER membrane (Aviram & Schuldiner, 2017) and implies recognition of the signal sequence by the signal recognition particle for delivery to the translocon (Gilmore, Blobel, & Walter, 1982; Meyer, Krause, & Dobberstein, 1982). Additional concepts, however, have emerged that postulate translation independent mRNA recruitment by ER membrane associated RNA binding proteins (Singer-Kruger & Jansen, 2014). In this case transcript recruitment is mediated by *cis* elements present on the mRNA itself and *trans*-acting RNA binding proteins (Kraut-Cohen et al., 2013). Given that Bfr1 mainly interacts with polysomes associated with the ER membrane (Lang et al., 2001) and that Bfr1 interacting mRNAs are enriched for secretory proteins (Lapointe et al., 2015) we analyzed whether *PMT1* and *PMT2* transcript localization was affected in *bfr1*Δ cells by subcellular fractionation. To this end soluble and membrane fractions of total cell extracts from *bfr1*Δ cells were separated by ultracentrifugation and *PMT2* transcript levels were analyzed in both fractions (Fig. 6A, B). The calculated *PMT2* mRNA ratio of membrane to soluble fraction is approximately 1 for wild type cells indicating equal distribution of *PMT2* between both fractions. In *BFR1* deficient cells the *PMT2* mRNA ratio does not significantly change (Fig. 6B).

**Fig. 6.**
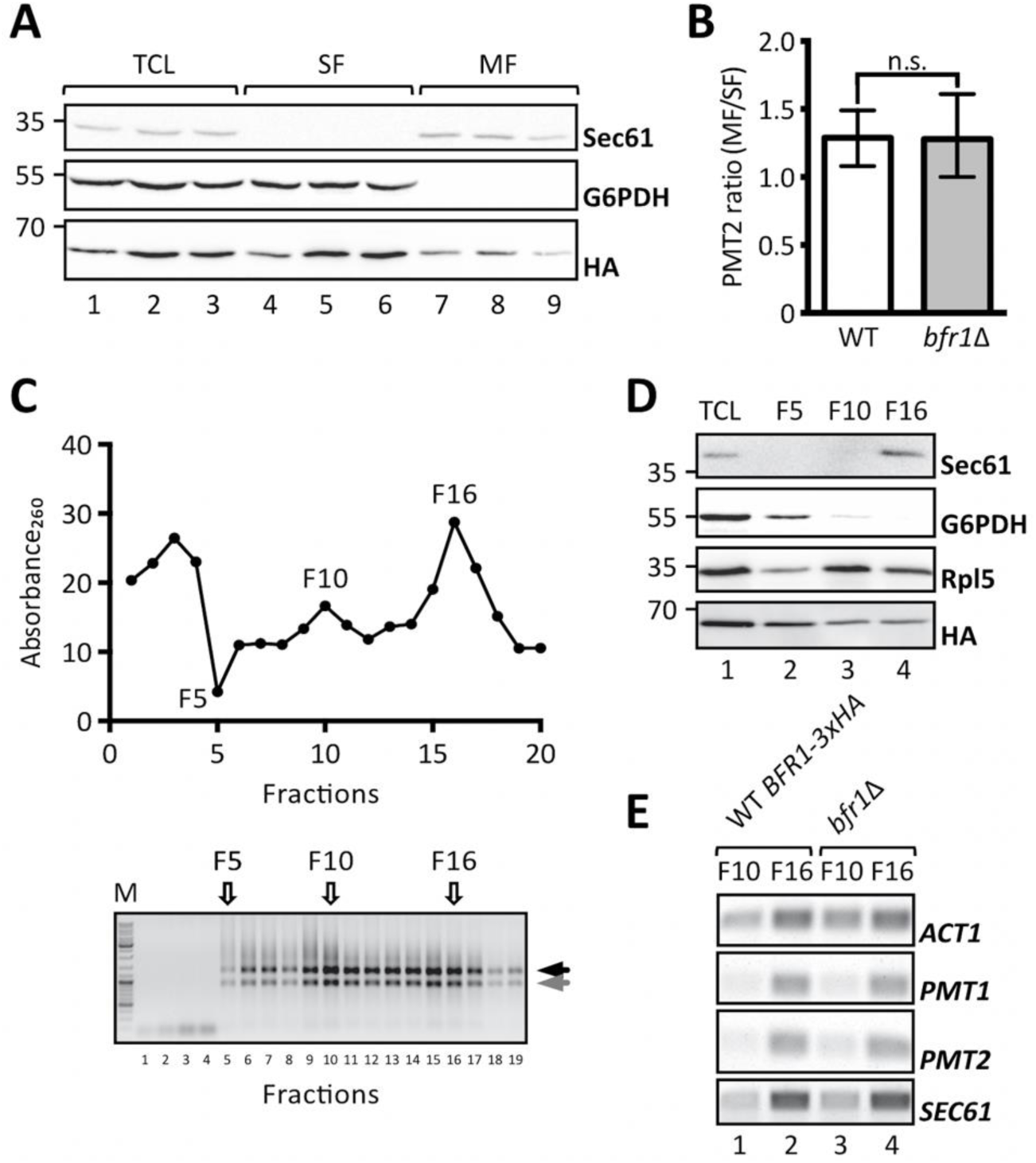
*BFR1* deletion does not affect *PMT1* and *PMT2* transcript localization. **(A)** and **(B)**: JCY017 (wild type BFR1-3xHA) cells were grown in YPD medium, lysed and total cell extracts were subjected to one step ultracentrifugation. **A)** Western blot analysis of total cell lysates (TCL), soluble and membrane fractions (SF and MF respectively) upon one step ultracentrifugation. Equivalents to 0.25 OD600 were resolved on a 12% PAA gel and detection was performed with the indicated antibodies. Sec61 served as a membrane marker and G6PDH as a cytosolic marker. Bfr1-3xHA was detected using the HA-tag. **B)** RT-PCR analysis of *PMT2* mRNA from soluble and membrane fractions upon one step ultracentrifugation. Total RNA was extracted from respective fractions and cDNA was prepared. *PMT2* mRNA from each fraction was normalized to *ACT1* mRNA. Results show the average membrane to soluble *PMT2* mRNA ratio from three independent experiments and error bars represent the confidence interval. For statistical significance one-sample t-test was performed on log2^−ΔΔCt^. **(C)**, **(D)** and **(E)**: JCY017 (wild type BFR1-3xHA) and *bfr1*Δ cells were grown in YPD medium, lysed and total cell extracts were subjected to sucrose step gradient centrifugation. **C)** Absorbance260 profile of fractions collected upon sucrose step gradient centrifugation (upper panel) and agarose gel electrophoresis of equivalent amounts of each fraction (lower panel). F5, F10 and F16 indicate fractions selected for further analysis. Black and grey arrows next to the agarose gel depict ribosomal subunits 60S and 40S. **D)** Western blot analysis of total cell lysates (TCL) and selected sucrose gradient fractions from JCY017 (wild type BFR1-3xHA) cells. 0.25 OD600 units of total cell extract and equivalents of selected fractions were resolved on a 12% PAA gel and detection was performed with the indicated antibodies. Sec61 and G6PDH were detected exclusively in fractions F16 and F5 respectively confirming successful fractionation. The ribosomal protein Rpl5 was mainly detected in fractions F10 and F16 that represent cytoplasmic and membrane bound polysomes respectively. The weaker Rpl5 signal detected in fraction F5 probably emanates from free cytosolic ribosomes. **E)** Semi-quantitative PCR analysis of *PMT1* and *PMT2* mRNA from sucrose gradient fractions F10 and F16 from JCY017 (wild type BFR1-3xHA) and *bfr1*Δ cells. Total RNA was extracted from respective fractions, cDNA was prepared and a 1:20 dilution was used as template in a standard DreamTaq PCR reaction. ACT1 that served as a loading control also shows strong engagement in the ER membrane containing fraction F16 in line with reports that the ER is a general translation hub even for cytosolic proteins (Pyhtila et al., 2008). Results are representative of two independent fractionations. N.s.=not significant

In addition, total cell extracts from wild type cells expressing fully functional HA-tagged Bfr1 and *bfr1*Δ cells were fractionated on a sucrose step gradient. Analysis of the RNA content of 20 collected fractions showed enrichment of ribosomes in fractions F10 and F16 (Fig. 6C). The respective control experiment was performed with EDTA supplemented lysates and resulted in the shift of both Absorbance260 peaks observed for F10 and F16 to soluble fractions in line with ribosomal disassembly (Suppl. Fig. 5). Analysis of specific marker proteins within F5, F10 and F16 reveals efficient separation of cytosolic and membrane fractions (Fig. 6D, compare lanes 2 and 4). All analyzed fractions contain ribosomes as assessed by the ribosomal protein Rpl5, however, to different extents. F10 represents the cytoplasmic polyribosome fraction whereas F16 contains ER membrane bound polysomes (Fig. 6D, compare Sec61 in lanes 3 and 4). Bfr1 was found throughout fractions consistent with reports of this cytosolic protein being associated with polyribosomes. Analysis of mRNA content in ribosome containing fractions F10 and F16 in wild type cells showed strong engagement of *PMT1*, *PMT2* and *SEC61* mRNA with ER membrane associated ribosomes whereas only minor amounts of these mRNAs were detectable in the cytoplasmic fraction F10 (Fig. 6E, lanes 1 and 2 respectively). In *BFR1* deficient cells distribution of neither *PMT1* and *PMT2* mRNAs nor *SEC61* and *ACT1* mRNAs was changed compared to wild type cells. In combination our data show that *PMT2* transcripts are equally distributed between the cytosolic and ER membrane bound polysomal fraction and that *PMT1* and *PMT2* mRNAs preferentially colocalize with membrane bound polyribosomes irrespective of Bfr1 presence.

### Bfr1 affects Pmt1 and Pmt2 translation

Next, we analyzed translation dynamics in wild type versus *bfr1*Δ cells by ribosome profiling, which provides a quantitative and high-resolution profile of *in vivo* translation and is based on deep sequencing of ribosome protected mRNA fragments (Ingolia, Ghaemmaghami, Newman, & Weissman, 2009). Protein synthesis rates are derived from average ribosomal density along mRNAs based on two fundamental assumptions: that all ribosomes complete translation and that elongation rates are similar among different mRNAs (Brar & Weissman, 2015). Ribosomal densities along transcripts show active translation and provide a snapshot of protein synthesis within the cell independent of transcript levels.

Ribosome profiling was performed in duplicate for both wild type and *BFR1* deficient cells. Replicates showed high correlation of reads per million mapped reads (RPM) values (r^2^=0.99 and r^2^=0.97 for wild type and *bfr1*Δ cells, respectively) (Suppl. Fig. 6A; Suppl. Table S2). RPM values of wild type and *bfr1*Δ cells also showed high correlation (r^2^=0.97) (Suppl. Fig. 6B; Suppl. Table S2) ruling out a generalized effect on translation. Statistical analysis revealed comparable subsets of genes significantly up- or downregulated at 0.01 false discovery rate (FDR) (red dots on Fig. 7A). For Pmt1 and Pmt2 ribosome profiling data demonstrate a *bfr1*Δ to wild type ratio of averaged RPMs of 0.581 and 0.596 respectively that corresponds to a significant 1.7-fold decrease of ribosomal footprint density and therefore active protein synthesis in *BFR1* deficient cells. This decrease in active translation correlates with the approximate 2-fold decrease in PMT protein abundance detected in *bfr1*Δ cells (Fig. 3A, B). In line with this observation, active translation of representative Bfr1 targets whose expression levels did not change upon *BFR1* deletion (Fig. 3C), remain unaffected with the exception of Kar2 (wild type/*bfr1*Δ=0.582) (Suppl. Table S2). Since *PMT1* and *PMT2* transcript levels do not change between wild type and mutant cells whereas ribosomal density is 1.7-fold lower these results reveal Bfr1 as a translational enhancer of Pmt1 and Pmt2. Furthermore, we combined our ribosomal footprint data with the Bfr1 mRNA interactome unraveled by Lapointe *et al*. (Lapointe et al., 2015). The 174 strongest mRNA interactors (Fig. 7B, class A) include 104 mRNAs encoding for proteins of the secretome (filled dots) defined by Ast *et al*. (Ast, Cohen, & Schuldiner, 2013). Translation of 35 mRNAs, all encoding secretome proteins, is significantly reduced in absence of *BFR1*, suggesting that Bfr1 preferentially affects translation of ER-targeted proteins. Intriguingly, GO functional annotation clustering identified among those targets protein glycosylation (*PMT1*, *OST1*, *PMT2*, *PMT3*, *PMT4*, *KTR1*, *STT3*, *ALG12*) and ergosterol biosynthesis (*ERG24*, *ERG3*, *NCP1*, *ERG4*, *ERG11*) as major functional clusters, pointing to Bfr1 as an important factor governing these processes.

**Fig. 7.**
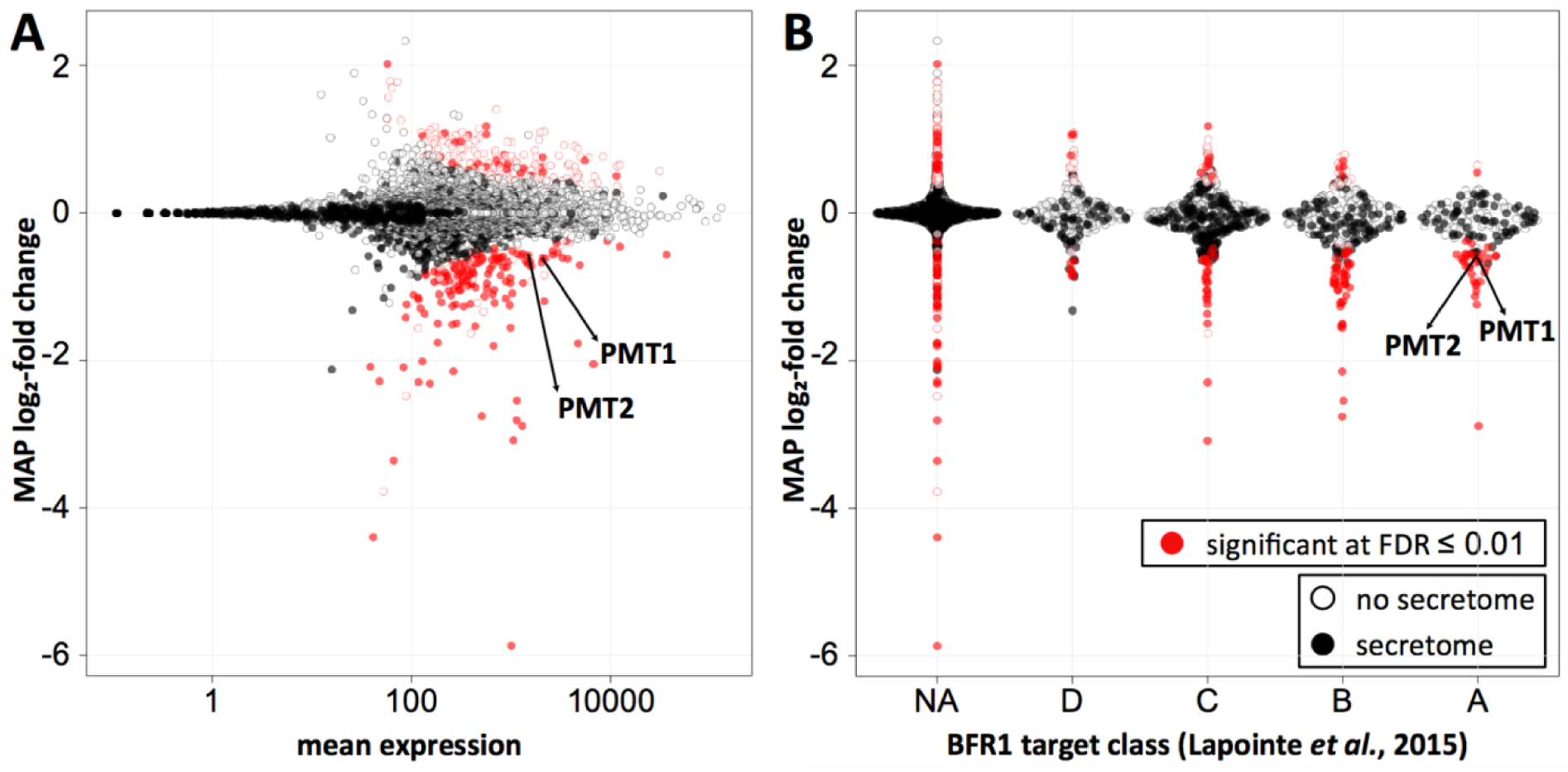
Bfr1 significantly enhances active translation of Pmt1 and Pmt2 and of a subgroup of secretory pathway proteins whose transcripts are strong Bfr1 interactors. **A)** MA plot showing active translation in *bfr1*Δ compared to wild type. Proteins for which translation is significantly affected in *bfr1*Δ versus wild type cells are depicted in red with filled red dots representing proteins assigned to the secretory pathway according to (Ast et al., 2013). Pmt1 and Pmt2 translation is significantly reduced in *bfr1*Δ cells. **B)** Ribosome profiling data were combined with data from an *in vivo* RNA tagging approach performed for Bfr1 (Lapointe et al., 2015). Classes A-D outlined in (Lapointe et al., 2015) contain candidates that are bound by Bfr1 to different extents: The strongest binders are in class A. In classes A and B most significantly affected proteins are down-rather than upregulated and are assigned to the secretory pathway (Ast et al., 2013). Pmt1 and Pmt2 are found among significantly downregulated proteins in class A. In **(A)** and **(B)** log2-fold changes were shrunken towards 0 using a Cauchy prior (Zhu, Ibrahim, & Love, 2019), the mode of the posterior distribution is shown. The amount of shrinkage is proportional to the gene-specific variance. FDR=false discovery rate, NA=not assigned

## DISCUSSION

In recent years, protein O-mannosylation proved to be critically important for ER protein quality control. O-mannosylation affects ER protein homeostasis at different levels. On one hand, stress sensors as well as other crucial components of protein folding and quality control machineries carry O-mannosyl glycans which may directly impact on their function (Castells-Ballester et al., 2018; Neubert et al., 2016). On the other hand, un- or misfolded proteins receive O-mannosyl glycans which label them for ER clearance (Xu & Ng, 2015b). In a first effort to identify factors that affect UPOM, the Pmt1-Pmt2 complex proved to be a central hub for ER protein quality control.

Among our screening hits we find several mutants that probably impact on ER protein folding but do not directly affect O-mannosylation of the UPOM-reporter (Suppl. Table S1). An example is *INO2* that encodes for a transcription factor responsible for derepression of phospholipid biosynthetic genes (Ambroziak & Henry, 1994). Membrane phospholipid perturbations have been linked to chronic ER stress in *S. cerevisiae* (Shyu et al., 2019). The presence of *INO2* as well as *SPF1* that was reported to cause ergosterol deficiency in the ER (Sorensen et al., 2019) further emphasizes the importance of ER membrane integrity to maintain the ER as a robust folding compartment in general. Most of the candidates, however, are linked to protein quality control as components of stress related pathways such as *sln1*Δ, *ptc1*Δ and *sic1*Δ that encode functional components of the HOG pathway as well as *oca1*Δ and *oca2*Δ that are involved in oxidative stress. Basal activity of the HOG pathway was shown to contribute to UPR induced accumulation of glycerol and thereby mediates resistance towards the ER stress inducing agent tunicamycin in *S. cerevisiae* (Torres-Quiroz, Garcia-Marques, Coria, Randez-Gil, & Prieto, 2010). Osmolytes such as glycerol are often referred to as “chemical chaperones” and have been shown to increase protein stability and restore ER homeostasis (Burg & Ferraris, 2008). Increased fluorescence of ER-GFP in *oca1*Δ and *oca2*Δ mutants might be explained by the recent finding that yeast UPR is inhibited by oxidative stress (Guerra-Moreno, Ang, Welsch, Jochem, & Hanna, 2019). With important components of the oxidative stress response missing yeast UPR could be more efficient in folding of the UPOM-reporter. However, general activation of UPR such as in *erj5*Δ (Carla Fama et al., 2007) and *erv25*Δ (Copic et al., 2009) or *hrd1*Δ mutants where ERAD is affected (Bays, Gardner, Seelig, Joazeiro, & Hampton, 2001), do not impact on ER-GFP folding (Xu et al., 2013) (Suppl. Table S1), suggesting a more specific role of stress related UPR for proper reporter folding.

Our screening further revealed unexpected links between ER-GFP, *per se* a non N-glycosylated protein, and N-glycosylation such as *cwh41*Δ and *ost3*Δ (Suppl. Table S1). *CWH41* encodes for α-glucosidase I that is responsible for trimming of the outermost glucose of N-glycans within the calnexin-calreticulin cycle thereby creating a time window before Mns1 and Htm1 mannosidases target the protein for degradation (Kostova & Wolf, 2003). Ost3 is one out of nine subunits of the yeast OST complex that together with Ost6 determines functionally distinct OST complexes (Schwarz, Knauer, & Lehle, 2005). Ost3 was recently reported to be necessary for N-glycosylation of Pmt2 (Zatorska et al., 2017) but no direct evidence of impaired Pmt2 enzymatic activity was obtained *in vivo*. However, ER-GFP oligomers that are indicative of ER-GFP misfolding (Xu et al., 2013) were significantly reduced in *ost3Δ* cells suggesting more efficient folding in the absence of Ost3 (Suppl. Fig. 2C).

In addition to Pmt1 and Pmt2, the strongest factors identified in the screen directly affecting O-mannosylation of ER-GFP are Psa1 and Bfr1 (Fig. 2B).

Psa1 catalyzes biosynthesis of GDP-mannose, the common sugar donor for Dol-P-Man production. Intriguingly, a second enzyme that contributes to GDP-mannose synthesis, the glucose-6-phosphate isomerase Pgi1 (Suppl. Fig. 3B), is found at immediate proximity to the screening threshold (Suppl. Table S1) suggesting that GDP-mannose availability might indeed be important for PMT activity. That carbohydrate donor levels affect PMT activity has been also suggested in studies performed in *S. cerevisiae* (Janik et al., 2003) and *Trichoderma reesei* (Zakrzewska et al., 2003) in which manipulation of GDP-mannose levels affects glycosylation. These preliminary data suggest a so far unknown link between carbohydrate metabolism and UPOM.

Bfr1 regulates Pmt1 and Pmt2 translation and therefore impacts on UPOM. Bfr1 is a cytoplasmic protein without any common RNA interacting motifs that was described as a component of polyribosome associated mRNP complexes in *S. cerevisiae* (Lang et al., 2001). Further, Bfr1 mediates localization of certain mRNAs to P-bodies (Simpson et al., 2014) and prevents P-body formation under normal conditions (Weidner et al., 2014) further supporting a function for Bfr1 in mRNA metabolism. P-bodies are dynamic ribonucleoprotein complexes where mRNA storage, translational repression or degradation occurs (Luo, Na, & Slavoff, 2018). Recent RNA binding studies that imply the presence of far more RNA binding domains than known to date (Albihlal & Gerber, 2018) in combination with multiple approaches that identify hundreds of different mRNAs interacting with Bfr1 (Hogan et al., 2008; Lapointe et al., 2015; Mitchell, Jain, She, & Parker, 2013) suggest a role for Bfr1 as an RNA binding protein and translational regulator itself.

In addition to Pmt1 and Pmt2, Bfr1 significantly affects active translation of all PMTs and of additional 322 genes from which nearly half show reduced translation in absence of Bfr1 (Fig. 7A; Suppl. Table S2). Among those we find the sterol reductase Erg4 that catalyzes the final step in ergosterol biosynthesis (Zweytick, Hrastnik, Kohlwein, & Daum, 2000) and that was recently described to be translationally regulated by Bfr1 (Manchalu, Mittal, Spang, & Jansen, 2019). We combined our data with Bfr1 interacting transcripts from Lapointe *et al*. (Lapointe et al., 2015) who reported Bfr1 targets to be highly enriched for mRNAs translated at the ER. In this “RNA Tagging” approach, Bfr1 interacting mRNAs were tagged with varying numbers of uridines by the poly(U) polymerase fused to Bfr1 depending on the strength of the interaction. Targets were classified into four groups based on the number of targeted RNAs and the length of the U-tag (class A encloses the strongest interactors). Crossing these datasets shows that Bfr1 controlled targets are enriched in classes A and B, which contain the strongest and most reliable Bfr1 binders. Among class A secretory proteins are Pmt1-4 and Erg4, as well as the OST subunits Ost1, Ost5 and Stt3 that form one out of two subcomplexes during OST complex assembly (Mueller et al., 2015). Given that these subcomplexes are intermediates that protect subunits from degradation they might play a decisive role in dynamics of OST complex formation and N-glycosylation. In addition, class A secretory proteins harbor several components of ergosterol biosynthesis (Erg3, Erg4, Erg11 and Erg24) and two iron homeostasis genes (Ftr1 and Smf3). This finding is particularly intriguing given the importance of iron for ergosterol biosynthesis and for Ire1 clustering and UPR activation (N. Cohen et al., 2017). A summary of all classified targets is available in Suppl. Table S2. Although a more detailed analysis of strong Bfr1 binders will be necessary to define the biological impact of Bfr1 mediated translation, our data strongly suggest a function of Bfr1 as a local translation factor at the ER membrane.

How does cytoplasmic Bfr1 regulate translation at the ER membrane? Our data strongly suggest that Bfr1 is not a prerequisite for PMT transcript recruitment to the ER, in agreement with similar observations for the Bfr1 target Erg4 (Manchalu et al., 2019). For Bfr1 this suggests two possible scenarios: Bfr1 could be targeted to the ER membrane via bound mRNAs as suggested for Erg4 (Manchalu et al., 2019) or Bfr1 could be associated with ER bound ribosomes before respective mRNAs reach the ER. It remains a challenging question for the future whether Bfr1 binds to mRNAs before or after their recruitment to the ER.

In a wider context our data together with transcriptomic data from others (Travers et al., 2000) reveal that ER stress is an important determinant of Pmt1-Pmt2 abundance (Fig. 3A, B; Fig. 4C) that is additionally controlled on a translational level by Bfr1 (Fig. 7). Interestingly Bfr1 is also a target of the UPR (Suppl. Fig. 7; (Travers et al., 2000)) suggesting that the function of Bfr1 is relevant to maintain protein homeostasis in the ER. Maximal Pmt1-Pmt2 expression depends on both, transcriptional activation of Pmt1-Pmt2 under cell stress conditions as well as elevated translation efficiency mediated by Bfr1. The fine-tuned coordination of Pmt1-Pmt2 protein abundance with ER stress further implies that O-mannosylation and protein folding must be balanced to ensure functionality of canonical target proteins and unfolded protein O-mannosylation, the latter being more sensitive to subtle changes of Pmt1-Pmt2 protein levels. Exactly adjusting Pmt1-Pmt2 activity to ER protein load most likely enables O-mannosylation of highly diverse protein substrates without unintentionally interfering with protein folding.

## MATERIALS AND METHODS

### Yeast Strains and Culture Conditions

*S. cerevisiae* strains used in this study are listed in Table 1. Strains derived from genetic libraries are underlined.

**Table 1.**
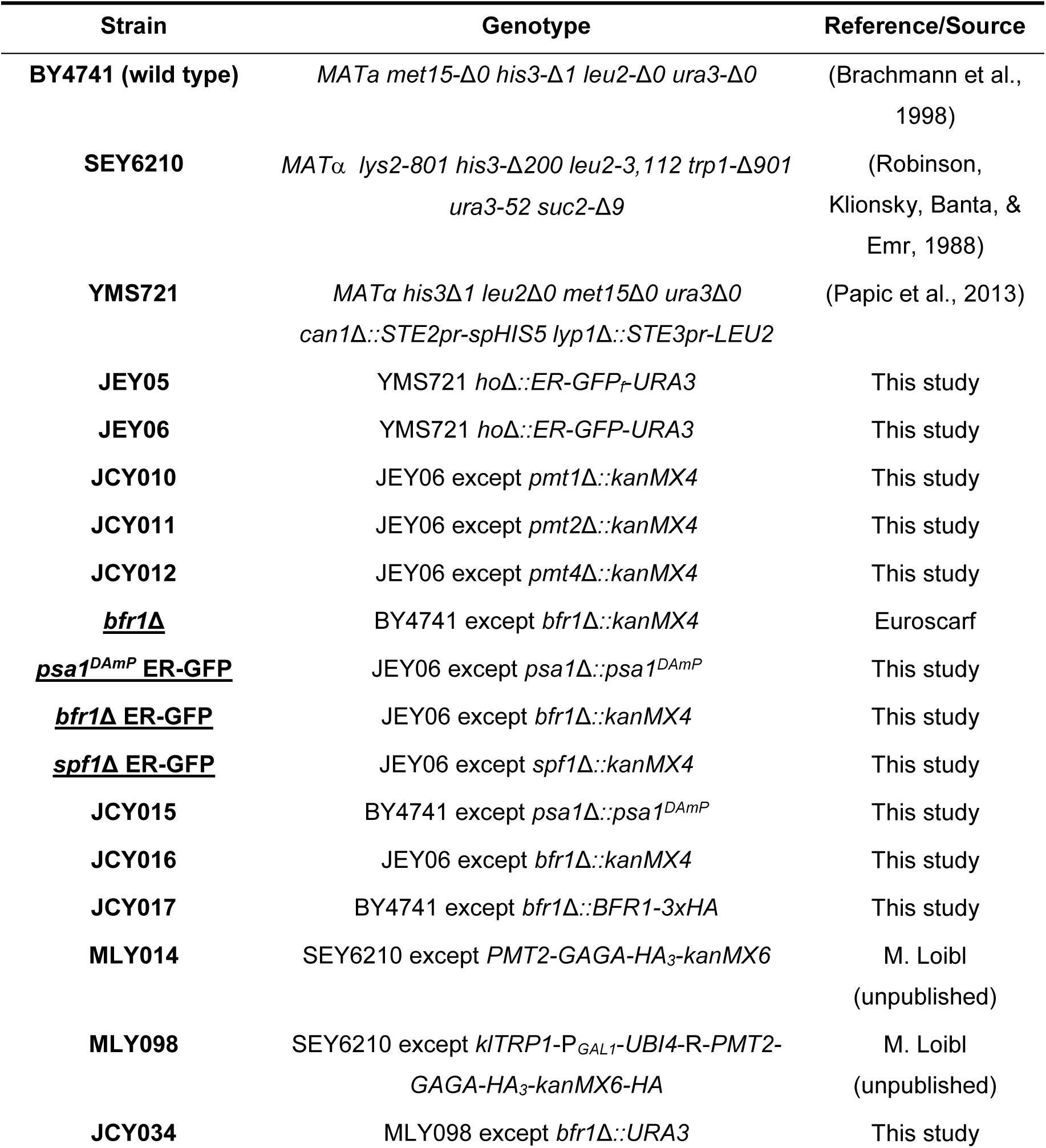
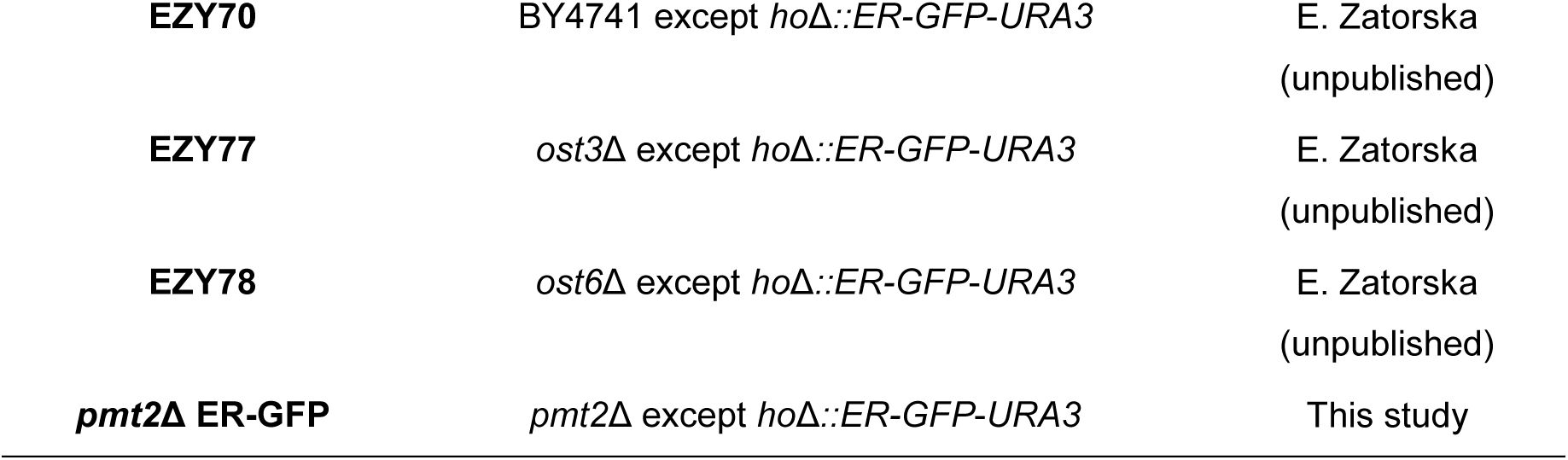
S. cerevisiae strains

Yeast cultures were grown in yeast extract-peptone-dextrose (YPD) or synthetic defined (SD) medium at 30°C. For auxotrophic selection corresponding amino acids were excluded from SD medium. For antibiotic-based selection cultures were supplemented with 400 µg/mL geneticin (#11811-031; Invitrogen; Waltham, MA, USA) or 100 mg/L nourseothricin (#96736-11-7; Werner BioAgents; Jena-Cospeda, Germany).

### Plasmids and Oligonucleotides

Plasmids used in this study are listed in Table 2. Sequences of oligonucleotides are available on request.

**Table 2.**
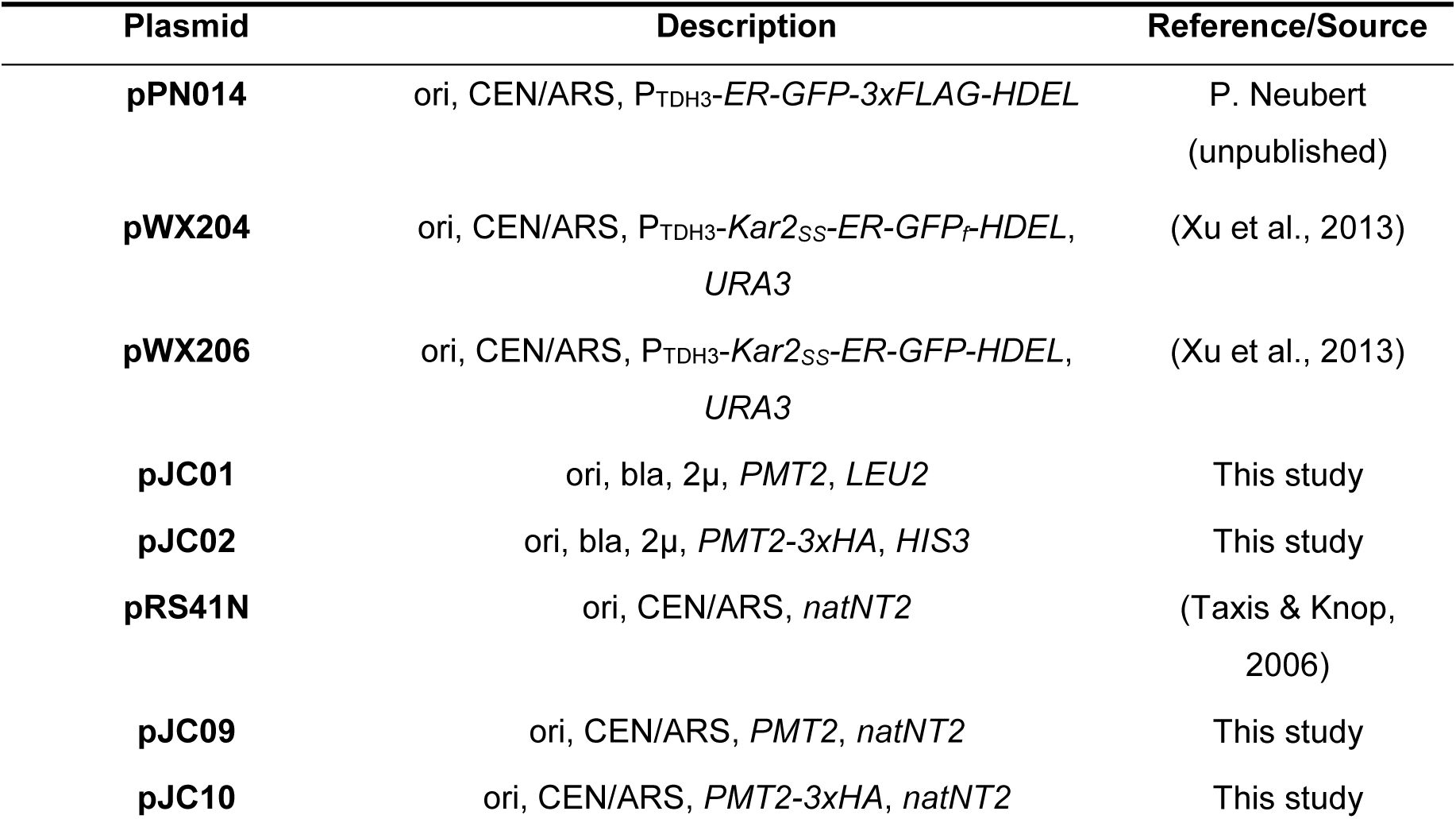
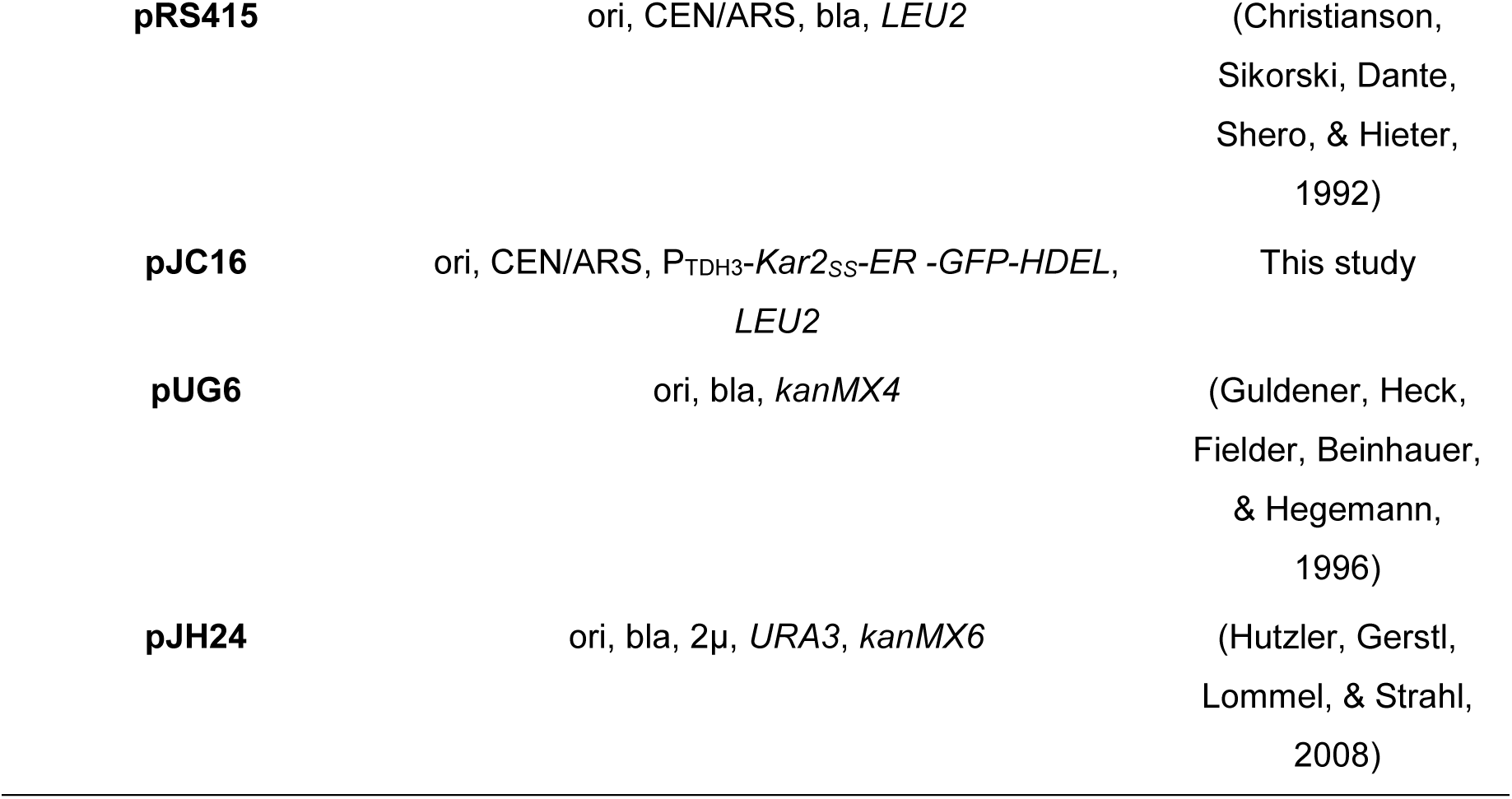
Plasmids

To construct plasmid pJC09 for Pmt2 expression the *SalI/PstI PMT2* fragment from pVG76 (Girrbach & Strahl, 2003) was cloned into pRS425 resulting in pJC01 and the *ApaI*/*SpeI PMT2* fragment from pJC01 was cloned into pRS41N. To construct plasmid pJC10 for Pmt2-3xHA expression the SalI/SmaI *PMT2-3xHA* fragment from pEZ43 was cloned into pRS423 resulting in pJC02 and the *ApaI*/*SpeI PMT2-3xHA* fragment from pJC02 was cloned into pRS41N. To construct pJC16 for Kar2SS-ER-GFP-HDEL expression the NotI/SalI *Kar2SS-ER-GFP-HDEL* fragment from pWX206 was subcloned into pRS415.

### ER-GFP Screening

#### Automated Library Generation

Query strain JEY06 expressing ER-GFP was constructed on the synthetic genetic array compatible strain YMS721 (Papic et al., 2013) and was integrated into yeast deletion (Giaever et al., 2002) and DAmP libraries (Breslow et al., 2008) following synthetic genetic array methodology (Y. Cohen & Schuldiner, 2011; Tong & Boone, 2006). Mating was performed on 1536-colony format YPD plates using a RoToR bench top colony arrayer (Singer Instruments; Somerset, UK). Resulting diploids were selected for deletion/DAmP libraries and ER-GFP markers *KanR* and *URA3*, respectively. Sporulation was induced by transferring cells to nitrogen starvation media for seven days and haploid cells were selected in histidine deficient SD plates to select for spores with an A mating type using canavanine and thialysine (Sigma-Aldrich) against remaining diploids alongside with previously mentioned selection markers.

#### High-throughput Microscopy

Microscopy screening was performed using an automated fluorescence microscopy setup as previously described (Breker, Gymrek, & Schuldiner, 2013). Cells were transferred from agar plates into liquid 384-well polystyrene growth plates using the RoToR arrayer. Liquid cultures were grown over night at 30°C in SD medium in a shacking incubator (LiCONiC Instruments; Liechtenstein). A JANUS liquid handler (PerkinElmer; Waltham, MA, USA) connected to the incubator was used to dilute the strains to an OD600 of approximately 0.2 into plates containing the same medium. Plates were incubated at 30°C for 4 h for cells to reach the logarithmic growth phase. Cultures were then transferred by the liquid handler into glass-bottom 384-well microscope plates (Matrical Bioscience; Spokane, WA, USA) coated with Concanavalin A (Sigma-Aldrich). After 20 minutes, wells were washed twice with SD-Riboflavin complete medium to remove non-adherent cells and to obtain a cell monolayer. Plates were then transferred to the ScanR automated inverted fluorescent microscope system (Olympus; Shinjuku, Japan) using a swap robotic arm (Hamilton; Bonaduz, Switzerland). Images of cells in 384-well plates were recorded in SD-Riboflavin complete medium at 24°C at GFP (excitation at 490/20 nm, emission at 535/50 nm) channel using a 60x air lens (NA=0.9) and with an ORCA-ER charge-coupled device camera (Hamamatsu; Shizuoka, Japan).

#### Image Analysis

Analysis of ER-GFP intensity was performed using the Olympus ScanR analysis software. Images were preprocessed by background subtraction and segmentation was done with the brightfield images and a series of shape conditions were applied as filters. The median GFP intensity for each strain was measured from the remaining objects for each strain. Dead cells appeared as high fluorescent outlier values and were removed with the ScanR software in an automated manner. Strains with insufficient number of detected objects (<25) as well as contaminated strains were removed from the analysis.

### Real-time Quantitative Polymerase Chain Reaction (RT-qPCR)

#### Total RNA Isolation

For total RNA isolation cells were grown to mid-log phase at 30°C. Ice-cold NaN3 was added to the culture to a final concentration of 100-200 mM and 5 OD600 units were harvested by centrifugation for 5 min at 3,000 g. Total RNA was isolated using the Universal RNA Purification Kit (Roboklon; Berlin, Germany) according to manufacturer’s instructions. For spheroplast generation prior to lysis lyticase (#L2524 Sigma-Aldrich Chemie; Munich, Germany) was added to the corresponding buffer. When indicated during the protocol RNase-free DNase (#M6101, Promega; Madison, WI, USA) was added to the RNA binding columns and incubated at RT for 10 min. For representative sets of samples RNA integrity was verified by agarose gel electrophoresis.

#### cDNA Synthesis

Two μg of total RNA were reverse transcribed into cDNA using the RevertAid First Strand cDNA Synthesis Kit (#K1622, Thermo Fisher Scientific, Bonn, Germany) with Oligo(dT)18 primers following manufacturer’s instructions.

#### Real-time Quantitative Polymerase Chain Reaction (RT-qPCR)

RT-qPCR was performed on the Rotor-Gene Q (Qiagen) using the qPCRBIO SyGreen Mix Lo-ROX (#PB20.11, PCR Biosystems, London, UK). PCR reactions were performed in a final volume of 12.5 μl containing × μl of 1:20 cDNA dilution and 0.4 mM of respective oligonucleotides. As technical replicates and for determination of RT-qPCR efficiency 1:100 and 1:1000 cDNA dilutions were included. Only RT-PCR reactions with efficiencies ranging from 0.9 to 1.1 were further analyzed. For calculation of either relative gene expression or fold-change in gene expression, both standard curve-based and 2-ΔΔCt methods were used. Statistical analysis was performed on three independent biological replicates. Statistical significance was assessed as individually stated.

### Preparation of Cell Extracts and Membranes

For cell extract preparation cells were grown to mid-log phase at 30°C. For end-point analyses or time course experiments ice-cold NaN3 was added to the culture to a final concentration of 100-200 mM and 10 or 20 OD600 units were harvested by centrifugation for 5 min at 3,000 g. Cells were washed and resuspended in 50 or 100 µl breaking buffer (50 mM Tris-HCl pH 7.4, 5 mM MgCl2) supplemented with protease inhibitors (1 mM PMSF, 1 mM benzamidine, 0.25 mM 1-chloro-3-tosylamido-7-amino-2-heptanone, 50 μg/mL l-1-tosylamido-2-phenylethyl chloromethyl ketone, 10 μg/mL antipain, 1 μg/mL leupeptin and 1 μg/mL pepstatin). Cell suspension was transferred to a tube with glass beads (ø 0.25-0.5 mm, #A553.1, Roth; Karlsruhe, Germany) and cells were subjected to mechanic lysis using the Hybaid RiboLyser (Thermo Fisher Scientific; Bonn, Germany) in four rounds of 25 s at 4.5 speed level. For cell extract preparation cell debris was pelleted by centrifugation for 5 min at 1,500 g. For membrane preparation total cell extracts were centrifuged for 1 h at 20,000 *g*. Membrane pellets were resuspended with a 0.3 mm syringe in equivalent volume of membrane buffer (20 mM Tris-HCl pH 8.0, 10 mM EDTA pH 8.0, 15% (v/v) glycerol) supplemented with protease inhibitors.

### Flag-tag Immunoprecipitation

For immunoprecipitation 20 OD600 units of cells grown to mid-log phase were subjected to membrane preparation with the following modifications: 1. Total cell extracts were centrifuged for 30 min at maximum speed (approximately 30,000 g). 2. Membrane buffer was supplemented with 1% Triton X-100 and samples were placed on a rotator mixer for 10 min at RT. 3. Samples were diluted 1:4 in TBS supplemented with 1mM PMSF and centrifuged for 15 min at 20,000 g to remove insoluble material. For Flag-tag immunoprecipitation samples were incubated with 100 µl of anti-FLAG M2 magnetic beads (#M8823, Sigma-Aldrich Chemie; Munich, Germany) for 4 h at 4°C. The bound fraction was eluted by addition of FLAG peptide to a final concentration of 0.3 µg/µl and further rotation for 1 h at 4°C. Demannosylation of ER-GFP was performed with α1-2,3,6 mannosidase (#9025-42-7, Sigma-Aldrich Chemie; Munich, Germany) according to manufacturer’s instructions.

### Cycloheximide Chase Experiments

Cells were grown under standard conditions in YPD medium and were initially sampled for time point zero. Cycloheximide was immediately added to a final concentration of 100 to 200 µM and equal amounts of cells were sampled at indicated time points. Sampled cells were treated with NaN3 to a final concentration of 20 mM to stop the chase and were kept on ice until the last sample was collected. Total cell extracts were analyzed via Western blot.

### Western Blot Analysis

Protein samples were denatured in 1x SDS-sample buffer for 10 min at 70°C and resolved in 12% sodium dodecyl sulfate polyacrylamide (SDS PAA) gels. Proteins were transferred to nitrocellulose membranes and visualized by enhanced chemiluminescence using ECL Prime Western Blotting Detection Reagent (#RPN2232, GE Healthcare; Chicago, IL, USA) and imager ImageQuant LAS 4000 (GE Healthcare; Chicago, IL, USA). Primary and secondary antibodies used in this study are summarized in Table 3.

**Table 3.** Antibodies

| Name | Description | Reference/Source |
| --- | --- | --- |
| αPmt1 | rabbit; 1:2,500 | (Strahl-Bolsinger & Tanner, 1991) |
| αPmt2 | rabbit; 1:2,500 | (Gentzsch, Immervoll, & Tanner, 1995) |
| αPmt4 | rabbit; 1:250 | (Girrbach & Strahl, 2003) |
| αSec61 | rabbit; 1:2,500 | (Stirling, Rothblatt, Hosobuchi, Deshaies, & Schekman, 1992) |
| αHA | mouse; 1:10,000 | #MMS-101R; Covance; Princeton, NJ, USA |
| αGas1 | rabbit; 1:2,500 | (Popolo, Grandori, Vai, Lacana, & Alberghina, 1988) |
| αWbp1 | rabbit; 1:2,500 | (te Heesen, Knauer, Lehle, & Aebi, 1993) |
| αKar2 | rabbit; 1:500 | |
| αOst3 | rabbit; 1:1,000 | Gift from M. Aebi |
| αG6PDH | rabbit; 1:5,000 | #A9521; Sigma-Aldrich Chemie; Munich, Germany |
| αGFP | rabbit; 1:2,500 | #A6455; Thermo Fisher Scientific; Waltham, MA, USA |
| αRpl5 | rabbit; 1:7,000 | Gift from E. Hurt |
| αmouse<sup>HRP</sup> | rabbit; 1:10,000 | #A9044; Sigma-Aldrich; Munich, Germany |
| αrabbit<sup>HRP</sup> | goat; 1:10,000 | #A6154; Sigma-Aldrich; Munich, Germany |

### Cell Fractionation coupled to RNA preparation

Both methods are adapted from (Aronov et al., 2015) and (Kraut-Cohen et al., 2013).

#### Cell Fractionation by One Step Ultracentrifugation

Cells grown to mid-log phase were treated with 100 µg/ml cycloheximide for 15 min before harvest. Cells equivalent to 20 OD600 units were harvested by centrifugation for 5 min at 3,000 g, washed with ice-cold SK buffer (1.2 M sorbitol, 0.1 M KPO4 pH 7.5, 100 μg/ml cycloheximide) and incubated for 5 min on ice. Cells were pelleted by centrifugation for 3 min at 500 g and were resuspended in 250 µl BRS buffer (50 mM Tris-HCl pH 7.6, 150 mM NaCl, 250 mM sorbitol, 30 mM MgCl2, 100 μg/ml cycloheximide, 200 U/ml RNasin ribonuclease inhibitor (#N2511, Promega; Madison, WI, USA)) supplemented with protease inhibitors as described for preparation of cell extracts and membranes. Mechanical lysis was performed with glass beads using the Hybaid RiboLyser in five rounds of 35 s at 4.5 speed level. For cell extract preparation cell debris was pelleted by centrifugation for 10 min at 1,000 g. 200 μl of cell extract were fractionated by ultracentrifugation at 48,000 g resulting in a soluble fraction and a membrane pellet. Membrane pellets were resuspended in 400-500 μl BMRS buffer (BRS buffer with 80 U/ml RNasin ribonuclease inhibitor) with a 0.3 mm syringe and ultracentrifugation was repeated. Total RNA was prepared from 100 µl of soluble and membrane fractions using the Universal RNA Purification Kit (Roboklon; Berlin, Germany) according to manufacturer’s instructions.

#### Cell Fractionation by Sucrose Step Gradient Centrifugation

Mid-log phase grown cells equivalent to 300 OD600 units were harvested by centrifugation for 5 min at 3,000 g, washed with ice-cold SK buffer and incubated for 5 min on ice. Cells were pelleted by centrifugation for 5 min at 2,500 g and were resuspended in 1.2 ml lysis buffer (10 mM Tris-HCl pH 7.5, 0.25 M sucrose, 30 mM MgCl2, 1 mM DTT, 100 μg/ml cycloheximide, 200 U/ml RNasin ribonuclease inhibitor) supplemented with protease inhibitors as described for preparation of cell extracts and membranes. Mechanical lysis was performed with glass beads using the Hybaid RiboLyser in four rounds of 45 s at 4.5 speed level. For cell extract preparation cell debris was pelleted by centrifugation for 10 min at 1,000 g. 900 μl of cell extract were diluted with lysis buffer to 2 ml final volume. For preparation of a discontinuous sucrose gradient 3 ml of a 1.5 M and 1.2 M sucrose buffer were added on top of a 2 M sucrose cushion. Total cell extract was loaded on top of the gradient and centrifugation was performed for 2.5 h at 232,000 g. The gradient was manually fractionated in 0.5 ml fractions and protein content of selected fractions was analyzed by SDS-PAGE. Total RNA was prepared from 300 µl of selected fractions using the Universal RNA Purification Kit (Roboklon; Berlin, Germany) according to manufacturer’s instructions. Semi-quantitative PCR was performed using the DreamTaq Green PCR master mix (#K1081, Thermo Fisher Scientific) according to manufacturer’s instructions. PCR was performed with 1 µl of a 1:20 dilution of cDNA in 23 or 25 cycles with a final primer concentration of 0.5 µM in a 20 µl reaction.

### Flow Cytometry

Cells expressing ER-GFP were grown to mid-log phase in the corresponding medium at 30°C. FACS analysis of 20,000 cells was performed using the cell analyzer BD FACSCanto™ (BD Biosciences; Heidelberg, Germany) in collaboration with the Flow Cytometry & FACS Core Facility (ZMBH, Heidelberg University; Heidelberg, Germany).

### Fluorescence Microscopy

Cells expressing ER-GFP were grown to mid-log phase in the corresponding medium at 30°C and microscopy was performed on standard glass plates using an LSM510-META confocal laser scanning microscope (Carl Zeiss; Jena, Germany) with x100 or x40 Plan Apochromat objectives. GFP signal (excitation 488 nm, Ar^+^ laser) was detected by using a bandpass emission filter for 505–530 nm.

### Ribosome Profiling

#### Sample Preparation

Wild type and *bfr1*Δ cells were grown to mid-log phase at 30°C and approximately 150 OD600 units were harvested using rapid filtration and flash freezing in liquid nitrogen. Frozen cell pellets were mixed with 750 µl of frozen lysis buffer droplets (20 mM Tris-HCl pH 8, 140 mM KCl, 10 mM MgCl2, 20% (v/v) NP-40, 100 μg/ml cycloheximide, 1x EDTA-free protease inhibitor cocktail (Roche; Basel, Switzerland), 0.02 U/μl DNase I (Roche; Basel, Switzerland), 40 μg/ml bestatin) and a metal ball in pre-chilled metal jars and lysed by mixer milling 2 min at 30 Hz (MM400, Retsch; Haan, Germany). Cell lysates were thawed in a 30°C water bath, transferred to low binding tubes and RNA concentration was determined by Nanodrop. Lysates were next subjected to RNase I digestion (10 U of RNase I per Abs260 unit) for 30 min at 4°C, the reaction was stopped by adding 10 µl of SUPERase-In RNase inhibitor (#LSAM2694, Invitrogen; Waltham, MA, USA) and lysates were cleared by 5 min centrifugation at 20,000 × gav.

Total ribosomes were collected by sucrose cushion centrifugation. Maximum of 400 µl of cleared lysate were loaded onto 800 µl of sucrose cushion buffer (20 mM Tris-HCl pH 8, 140 mM KCl, 10 mM MgCl2, 100 μg/ml cycloheximide, 1x EDTA-free protease inhibitor cocktail (Roche; Basel, Switzerland), 25% (v/v) sucrose) in sucrose cushion tubes and centrifuged for 90 min at 75,000 rpm and 4°C in a TLA120-rotor (Beckman; Indianapolis, IN, USA). Pellets were resuspended in lysis buffer by continuous agitation at 4°C and transferred to non-stick tubes.

#### Ribosome-protected Footprint mRNA Extraction

mRNA footprints were extracted from processed samples by phenol-chloroform extraction. In brief, ribosome pellets were brought to a final volume of 700 µl with lysis buffer and mixed with 40 µl 20% (v/v) SDS to precipitate the protein content. 750 µl of pre-warmed (65°C) acid phenol was added and samples were incubated for 5 min at 65°C and 1,400 rpm shaking; and chilled for 5 min on ice. Next, samples were centrifuged for 2 min at 20,000 × gav and the aqueous phase was transferred to a new tube. 700 µl of hot phenol were again added and samples were incubated 5 min at room temperature with occasional vortexing. 600 µl of chloroform were added and mixed by vortexing. Samples were centrifuged for 1 min at 20,000 × gav and the aqueous phase was transferred to a new tube. To precipitate nucleic acids, ∼ 650 µl of the sample were mixed with 1:9 equivalence volume of 3 M NaOAc pH 5.5, 1 equivalence volume of isopropanol and 2 µl of Glycoblue, mixed by vortexing and chilled overnight at −80°C.

Next, RNA samples were centrifuged for 2 h at 20,000 × gav and 4°C and the pellet was washed with 750 µl ice-cold 70% ethanol. Centrifugation was repeated for 2 min and the pellet was dried for 2 min at 65°C. Pellets were finally resuspended in 20-50 µl of 10 mM Tris-HCl pH 7.

RNA enrichment was verified by Bioanalyzer RNA Nanochip (Agilent) and total RNA concentration was determined by Nanodrop after diluting RNA samples in water and 10 mM Tris-HCl pH 7, respectively.

#### Deep Sequencing Library Preparation

Total translatome analysis was performed according to (Doring et al., 2017) with some modifications. RNA samples were heated at 80°C for 2 min and 40-50 mg of RNA were loaded onto 15% TBE-Urea polyacrylamide gels (Invitrogen; Waltham, MA, USA) in 1x TBE (Ambion) and run for 65 min at 200 V. Gels were stained for 20 min with SYBR gold (Invitrogen) and ribosome footprints were recovered from the gels by excising sections of 21 to 33 nucleotide size. Gel pieces were placed into 0.5 ml gel breaker tubes and centrifuged for 3 min at 20,000 × gav. Remaining pieces were transferred to a fresh 1.5 ml tube, resuspended with 10 mM Tris-HCl pH 7 and incubated for 15 min at 70°C in a thermomixer with maximum shaking. The gel slurry was then transferred to a Spin-X cellulose acetate column (#60702, Thermo Fisher Scientific; Waltham, MA, USA) and centrifuged for 3 min at 20,000 × gav. Flow through was transferred to a fresh pre-cooled non-stick tube on ice. Nucleic acids samples were precipitated as described in the latter section. Next, RNA samples were centrifuged for 2 h at 20,000 × gav and 4°C and the pellet was washed with 750 µl ice-cold 70% ethanol. Centrifugation was repeated for 2 min and the pellet was dried for 2 min at 65°C. Pellets were finally resuspended in 15 µl of 10 mM Tris-HCl pH 7 and transferred to a fresh non-stick tube. *To dephosphorylate 3’ ends of ribosome footprints*, a master mix was prepared containing 2 µl 10x T4 polynucleotide kinase buffer without ATP (NEB) and 1 µl murine RNase inhibitor per sample and 3 µl were added to each sample together with 2 µl truncated T4 polynucleotide kinase (#M0201, NEB; Frankfurt/Main, Germany). Samples were incubated for 2 h at 37°C and the enzyme was deactivated after the reaction by 10 min incubation at 75°C. At this point, nucleic acids were again precipitated as previously indicated. Samples were centrifuged for 1 h at 20,000 × gav and 4°C and RNA pellets were washed with 70% ethanol and resuspended in 15 µl of 10 mM Tris-HCl pH 7 and transferred to a fresh non-stick tube as previously indicated. RNA concentration was measured by Bioanalyzer RNA Nanochip (Agilent) and by nanodrop after diluting RNA samples in water and 10 mM Tris-HCl pH 7, respectively. *For 3’ L1 linker ligation*, samples were diluted to a final RNA concentration of 10 pmol in 10 µl of 10 mM Tris-HCl pH 7 and denatured for 2 min at 80°C. A master mix was prepared containing 16 µl 50% sterile filtered PEG MW 8000, 4 µl DMSO, 4 µl 10x T4 RNA Ligase 2 buffer and 2 µl murine RNase inhibitor. Master mix was added to each sample together with 1 µl truncated T4 RNA Ligase 2 (#M0239, NEB; Frankfurt/Main, Germany). Ligation was carried out for 2 h at 23°C and nucleic acids were precipitated, RNA pellets were washed with 70% ethanol as previously indicated and resuspended in 6 µl of 10 mM Tris-HCl pH 7. 3’-linked footprints were denatured at 80°C for 2 min and purified on 10% TBE-Urea polyacrylamide gels (Invitrogen) in 1x TBE (Ambion) run for 50 min at 200 V. Gels were stained for 20 min with SYBR gold (Invitrogen) and 3’-linked footprints were recovered from the gels by excising sections of 64 nucleotide size (footprint + L1). Similar to the previous in-gel purification, gel pieces were placed into 0.5 ml gel breaker tubes and centrifuged for 5 min at 20,000 × gav. Remaining pieces were transferred to a fresh 1.5 ml tube, resuspended with 10 mM Tris-HCl pH 7 and incubated for 15 min at 70°C in a thermomixer with maximum shaking. The gel slurry was then transferred to a Spin-X cellulose acetate column (#60702, Thermo Fisher Scientific; Waltham, MA, USA) and centrifuged for 3 min at 20,000 × gav. Flow through was transferred to a fresh pre-cooled non-stick tube on ice, nucleic acids were precipitated, RNA pellets washed with 70% ethanol as previously indicated and resuspended in 6 µl of 10 mM Tris-HCl pH 7.

*To generate ssDNA* 3’-linked footprint fragments were reverse transcribed. A master mix containing 1 µl 10 mM dNTP mix, 1 µl 25 µM Linker L1’L20 and 1.5 µl DEPC H2O was prepared and added to the samples. Samples were incubated for 5 min at 65°C and 4 ml 5x FSB buffer (Invitrogen), 1 ml murine RNase inhibitor, 1 ml 0.1 M DTT (Invitrogen) and 1 ml Superscript III (Invitrogen) were added. Reverse transcription was performed for 30 min at 50°C and the reaction was quenched by adding 2.3 ml 1 N NaOH and further incubating for 15 min at 95°C. Samples were denatured for 2 min at 70°C and run on a 10% TBE-Urea polyacrylamide gel for 70 min at 200 V. Gels were stained as described before, desired bands were excised and nucleic acids were extracted as mentioned earlier except remaining gel pieces were mixed with 0.5 ml 10 mM Tris-HCl pH 8. Nucleic acids were precipitated by adding 1:16 equivalence volume of 5 M NaCl and 1:500 equivalence volume of 0.5 M EDTA together with 1 equivalence volume of isopropanol and 2 µl of Glycoblue. Precipitation was performed at −20°C overnight and pellets were washed with 70% ethanol and resuspended in 15 µl 10 mM Tris-HCl pH 8 as previously described.

To circularize ssDNA a master mix containing 2 µl 10x CircLigase buffer, 1 µl 1 mM ATP, 1 µl 50 mM MnCl2 was added to the samples together with 1 µl CircLigase (EPICENTRE). Reaction was carried out for 1h at 60°C and the enzyme was inactivated by further incubation for 10 min at 80°C. 1 µl of circularized ssDNA was used as a template for 4 technical replicates of Phusion-based PCR using the following mix and PCR program: PCR mix (62.6 µl DEPC H20, 16.7 µl 5x HF buffer, 1.7 µl 10 mM dNTPs, 0.4 μl 100 mM barcoding primer, 0.4 µl 100 mM PCR primer L1’, 0.8 µl Phusion polymerase), PCR program (Initial denaturation: 98°C, 30s, (Denaturation; 98°C, 10s, Annealing: 60°C, 10s, Elongation: 72°C, 5s)x10 cycles). One tube was removed from the PCR reaction after cycles 7, 8, 9 and 10. Samples were run on a 8% TBE polyacrylamide gel (Invitrogen) in 1x TBE (Ambion) for 55 min at 180 V. Gels were stained as mentioned before, desired bands from each PCR reaction were excised and DNA was extracted as described before for the ssDNA samples. Size distribution of DNA fragments was determined by Bioanalyzer, concentration was determined by Qubit (#Q32852, Invitrogen) and samples were sequenced on a HiSeq (Illumina).

Sequenced reads were processed as described previously (Galmozzi, Merker, Friedrich, Doring, & Kramer, 2019) using standard analysis tools (Bowtie2, Tophat2) and python scripts adapted to *S. cerevisiae*. For each read, the P-site position was determined using a 5’ offset of 15 nucleotides. Only reads with a length of 25-35 nucleotides were used. Reads with P sites falling within an annotated ORF were counted, differential expression analysis was performed with DESeq2 (Love, Huber, & Anders, 2014) and false discovery rate was controlled using the Benjamini-Hochberg procedure (Benjamini & Hochberg, 1995) with independent hypothesis weighting (Ignatiadis, Klaus, Zaugg, & Huber, 2016).

## SUPPLEMENTAL TABLES

**Suppl. Table S1.** UPOM screening results

**Suppl. Table S2.** Ribosome profiling

## SUPPLEMENTAL FIGURES

**Suppl. Fig. 1.**
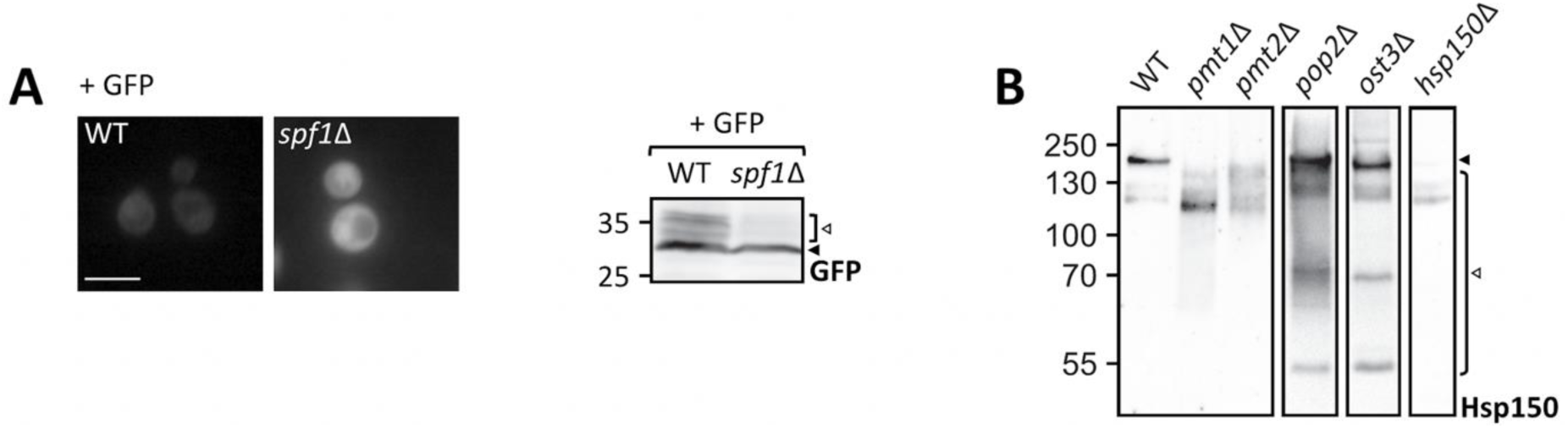
Evaluation of UPOM screen hit *spf1*Δ using ER-GFP and of *pop2*Δ and *ost3*Δ using Hsp150. **A)** Analysis of ER-GFP subcellular localization in wild type (BY4741) cells and in a screening independent *spf1*Δ strain and Western blot analysis of ER-GFP O-mannosylation in total cell extracts from the same strains. Cells were transformed with ER-GFP and grown in SD supplemented with uracil for selection before being imaged under standard conditions (scale bar 5 μm) or lysed for Western blot analysis. Equivalents to 0.2 OD600 were resolved on a 12% PAA gel and detection was performed with an anti-GFP antibody. **B)** Western blot analysis of Hsp150 in *pop2*Δ and *ost3*Δ cells used for clustering of UPOM screen hits. Viable single deletion mutants were retrieved from the Euroscarf collection and subjected to heat shock to induce Hsp150 secretion. Proteins of the medium were resolved on 8% PAA gels and detection was performed with an anti-Hsp150 antibody. Media from wild type and *hsp150*Δ cells were included as positive controls. Hsp150 fully glycosylated and hypoglycosylated fractions are indicated with black and white arrows respectively. Results from 100 deletion mutants identified as screening hits are summarized in Suppl. Table S1.

**Suppl. Fig. 2.**
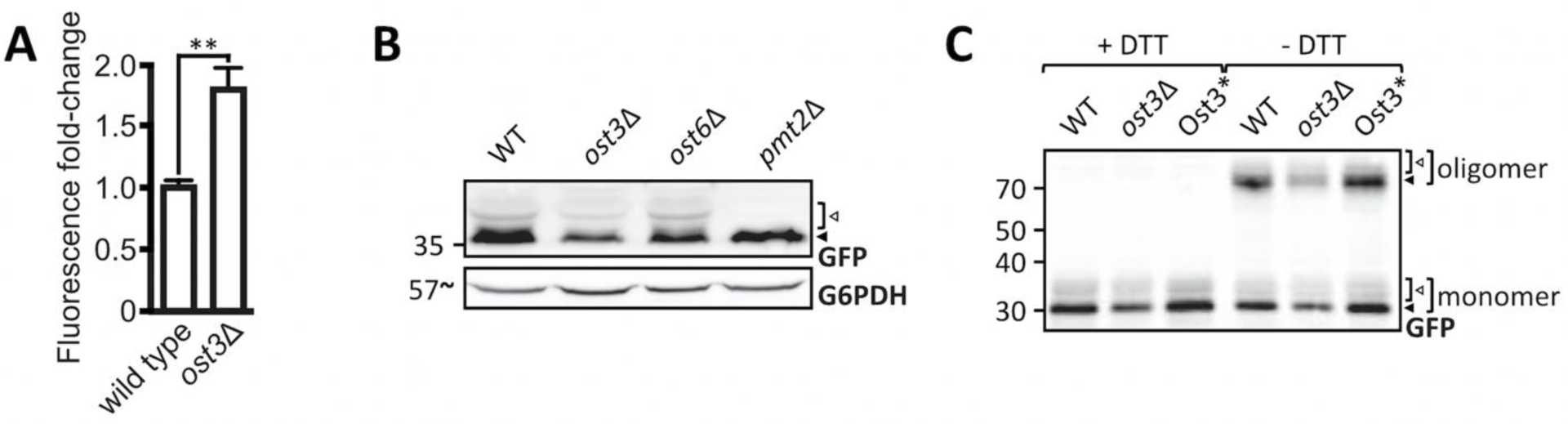
Evaluation of UPOM screen hit *ost3*Δ using ER-GFP. **A)** Flow cytometry analysis of EZY83 (wild type) and EZY82 (*ost3*Δ) cells grown to mid-log phase. Fluorescent signal resulting from analysis of 20000 cells was normalized to wild type and results are plotted as fold-change. Error bars represent the range of values from three independent experiments ± SD. For statistical significance Tukey’s HSD test was performed. Western blot analysis of **B)** ER-GFP O-mannosylation in total cell extracts from EZY70 (wild type), EZY77 (*ost3*Δ), EZY78 (*ost6*Δ) and *pmt2*Δ ER-GFP (*pmt2*Δ) strains and **C)** ER-GFP oligomerization in total cell extracts from EZY83 (wild type), EZY82 (*ost3*Δ) and EZY84 (Ost3*) strains. 20 μg of protein were resolved on a 12% PAA gel and detection was performed with an anti-GFP antibody. G6PDH was used as loading control. In **(C)** protein was denatured in sample buffer containing or lacking DTT. Monomeric and oligomeric ER-GFP as well as the main ER-GFP signal and higher O-mannosylated GFP-fractions are depicted by black and white arrows respectively.

**Suppl. Fig. 3.**
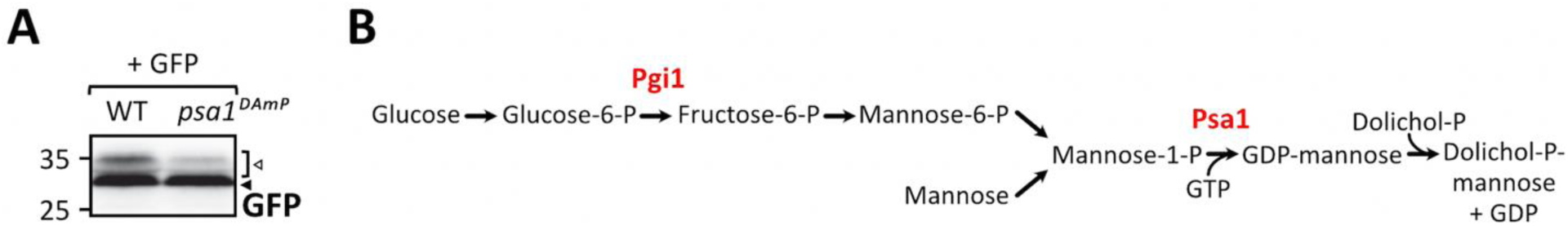
Evaluation of UPOM screen hit *psa1*Δ using ER-GFP. **A)** Western blot analysis of ER-GFP O-mannosylation in total cell extracts from wild type (BY4741) cells and the screening derived *Psa1^DAmP^* mutant. Equivalents to 0.2 OD600 were resolved on a 12% PAA gel and detection was performed with an anti-GFP antibody. Arrows on the right indicate the main GFP signal (black arrow) and signals emanating from higher O-mannosylated GFP fractions (white arrow). **B)** Scheme of cytosolic pathways producing GDP-mannose with the UPOM screen hits Pgi1 and Psa1 highlighted in red.

**Suppl. Fig. 4.**
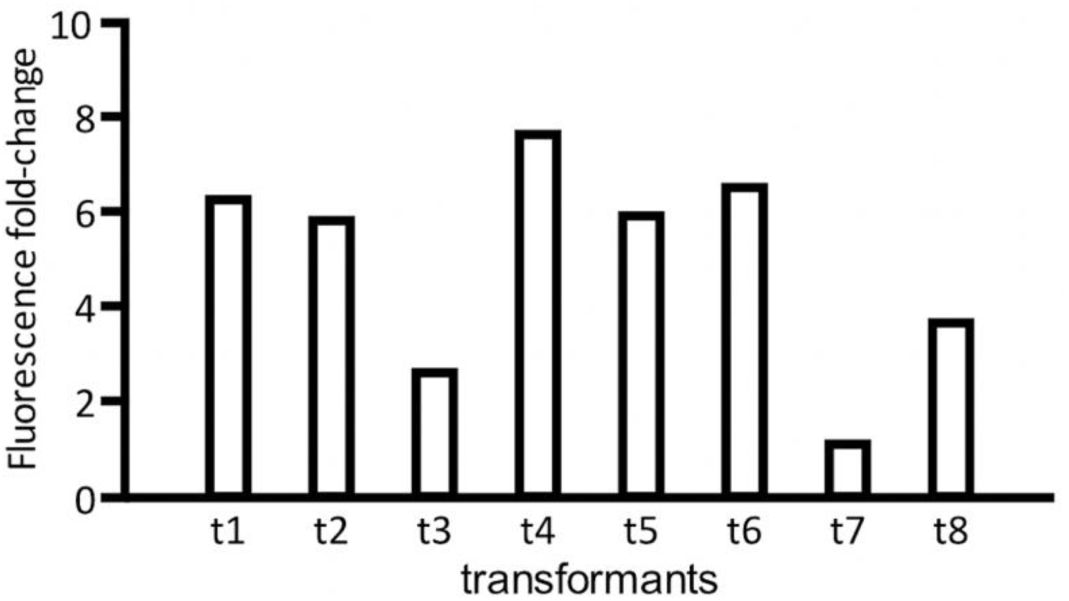
Analysis of screen-independent *bfr1*Δ knockout transformants. *BFR1* was knocked out in JEY06 (wild type ER-GFP) by homologous recombination. The knockout cassette containing up- and downstream *BFR1* homologous regions and *KanMX6* was generated via PCR from pUG6. After selection, *KanMX6* insertion was verified by PCR and eight independent transformants were grown in YPD and analyzed via flow cytometry. Fluorescent signal resulting from analysis of 20000 cells was normalized to wild type and results are plotted as fold-change.

**Suppl. Fig. 5.**
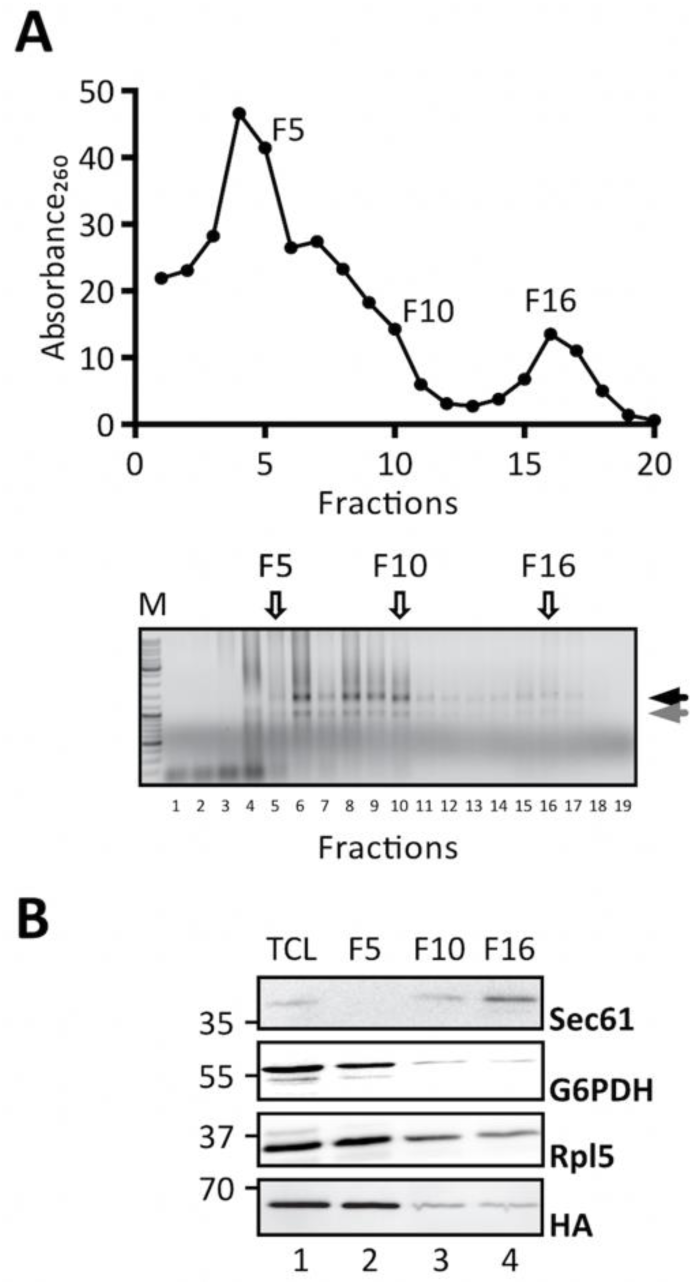
Control experiment to the *PMT1* and *PMT2* transcript localization experiment depicted in Fig. 6. The experiment was performed as described in Fig. 6, however, in presence of EDTA. EDTA was added to total cell lysates to a final concentration of 30 mM. **A)** EDTA treatment causes the rRNA associated Absorbance260 to shift from ribosome associated fractions F10 (free ribosomes) and F16 (ribosomes in polysomes) to the cytoplasmic fraction F5 (upper panel) as well as disassembly of ribosomal subunits depicted by the black and grey arrow (lower panel). **B)** In line with Bfr1 being primarily associated with ribosomes, EDTA treatment leads to redistribution of Bfr1 (HA-signal) from F10 and F16 to F5.

**Suppl. Fig. 6.**
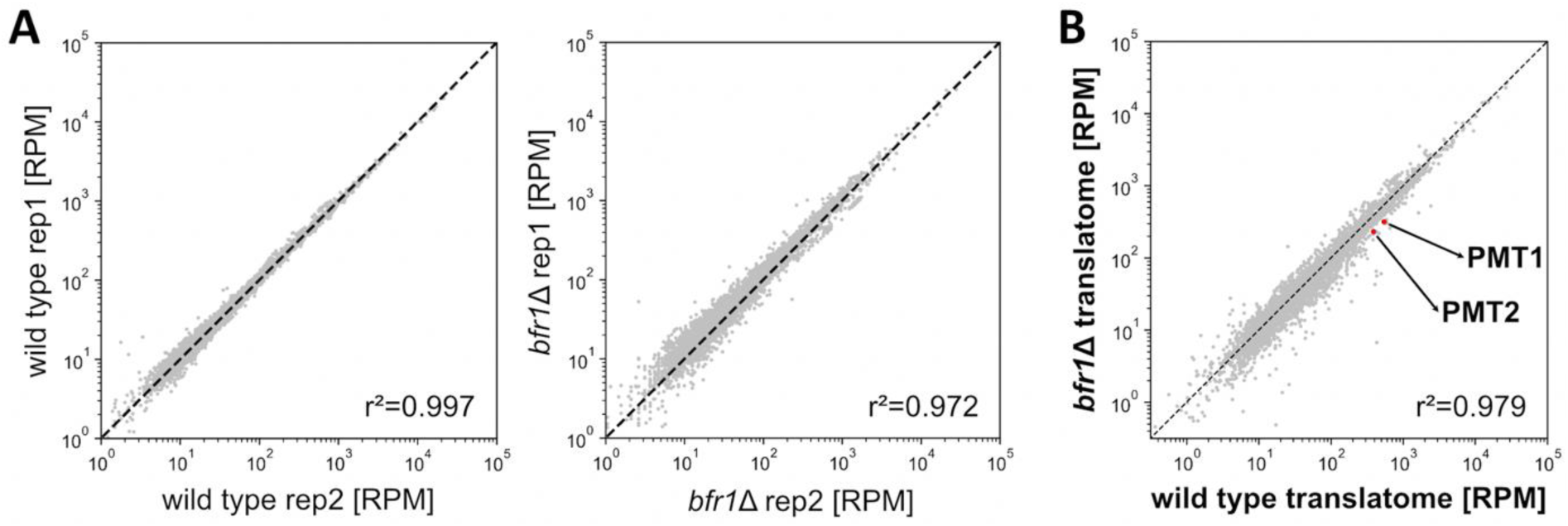
Active translation of Pmt1 and Pmt2 is significantly reduced in the absence of Bfr1. **A)** Correlation scatter plots of replicates (rep) from wild type and *bfr1*Δ cells. **B)** Scatter plot comparing normalized ribosome densities between wild type and *bfr1*Δ cells across the *S. cerevisiae* transcriptome. Pmt1 and Pmt2 are 1.7-fold downregulated in *bfr1*Δ versus wild type cells (indicated with red dots). Data were normalized to RPMs (reads per million mapped reads).

**Suppl. Fig. 7.**
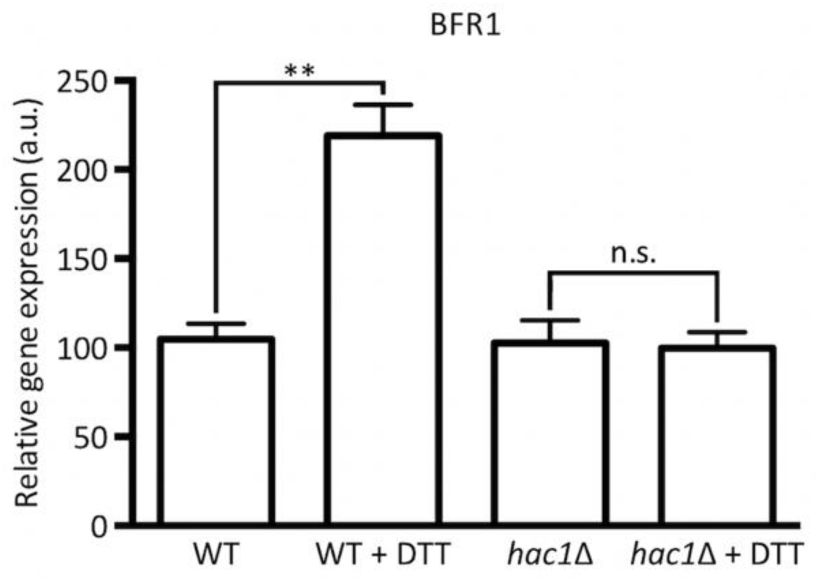
*BFR1* is induced by the UPR. RT-PCR analysis of *BFR1* mRNA levels in wild type and *hac1*Δ cells in response to DTT. Wild type (BY4741) and *hac1*Δ (Euroscarf) cells were treated with 2.2 mM DTT for 60 min, total RNA was extracted, and cDNA was prepared and used as a template for RT-PCR. Results show mRNA abundance with respect to *ACT1* mRNA from three independent experiments ± SD. For statistical significance a two-tailed t-test was applied (n=3). N.s.=not significant

## Author Contributions

Joan Castells-Ballester, Guenther Kramer, Maya Schuldiner and Sabine Strahl conceived of and designed the experiments. Joan Castells-Ballester, Lihi Gal, Ilgin Kotan, Daniela Bausewein and Ewa Zatorska performed the experiments. Joan Castells-Ballester, Natalie Rinis and Ilia Kats analyzed and evaluated data. Natalie Rinis, Joan Castells-Ballester and Sabine Strahl wrote the paper. Maya Schuldiner and Bernd Bukau edited the paper.

## Acknowledgments

We are grateful to Davis Ng for providing plasmids pWX204 and pWX206. We thank Anke Metschies and Silvia Chuartzman for excellent technical assistance, Sven Klassa for his help on the analysis of *psa1^DAmp^* and *pgi1^DAmp^* mutants and Jakob Engel for generating yeast strains JEY05 and JEY06.

